# The Structure of ApoB100 from Human Low-density Lipoprotein

**DOI:** 10.1101/2024.02.28.582555

**Authors:** Zachary T. Berndsen, C. Keith Cassidy

## Abstract

Low-density lipoprotein (LDL) plays a central role in lipid and cholesterol metabolism and is a key molecular agent involved in the development and progression of atherosclerosis, a leading cause of mortality worldwide. Apolipoprotein B100 (apoB100), one of the largest proteins in the genome, is the primary structural and functional component of LDL, yet its size and complex lipid associations have posed major challenges for structural studies. Here we overcome those challenges and present the first structure of apoB100 from human LDL using an integrative approach of cryo-electron microscopy, AlphaFold2, and molecular dynamics-based refinement. The structure consists of a large globular N-terminal domain that leads into a ∼58 nm long x 4 nm wide continuous amphipathic β-sheet that wraps completely around the circumference of the particle, holding it together like a belt. Distributed symmetrically across the two sides of the β-belt are 9 strategically located inserts that vary in size from ∼30-700 residues and appear to have diverse functions. The largest two form long flexible strings of paired amphipathic helices that extend across the lipid surface to provide additional structural support through specific long-range interactions. These results suggest a mechanism for how the various domains of apoB100 act in concert to maintain LDL shape and cohesion across a wide range of particle sizes. More generally, they advance our fundamental understanding of LDL form and function and will help accelerate the design of potential new therapeutics.

## Introduction

Low-density lipoprotein (LDL) is a complex macromolecular nanoparticle that facilitates the transport of fat-soluble molecules such as triglycerides and cholesterol between the liver and peripheral tissues [1]. LDL particles range in diameter from ∼17-28 nm and consist of a monolayer of phospholipid and free cholesterol (CHS) with embedded apolipoproteins, all surrounding a core of primarily triglycerides (TGs) and cholesteryl esters (CE) [1–3]. LDL particles are heterogeneous in size, shape, and composition, and multiple subclasses of LDL are categorized with different properties and metabolic origins [4]. Some start in the liver as large (∼40-70 nm) triglyceride-rich very low-density lipoprotein (VLDL), then evolve through successive metabolic stages—first into intermediate-density lipoprotein (IDL), then into the smaller triglyceride-depleted cholesterol-rich LDL [1]. Other subclasses are synthesized *de novo* and secreted directly as LDL [4]. Central to the structure and function of every LDL particle is a single copy of apolipoprotein B100 (apoB100), a ∼550KDa protein that not only governs the particle’s hepatic synthesis and structural cohesion but also mediates interactions with the LDL receptor (LDL-r), thereby directing cellular LDL uptake [1, 5, 6].

While instrumental in lipid and cholesterol homeostasis, LDL also has important pathological significance as elevated serum levels contribute to the development and progression of atherosclerosis, a leading cause of cardiovascular morbidity and mortality worldwide [7–9]. This pathogenic process initiates when LDL particles become retained in the arterial wall and oxidized. The uptake of oxidized LDL by macrophages transforms them into cholesterol-laden foam cells, marking the onset of plaque formation [8]. Therefore, therapeutic strategies against atherosclerosis have prominently focused on lowering LDL levels, with statins and proprotein convertase subtilisin/kexin type 9 (PCSK9) inhibitors emerging as the most efficacious pharmaceutical agents [10, 11]. In addition to dietary influences, there are >100 mutations in the APOB gene known to affect LDL levels, some associated with early-onset disease and mortality [12–14].

LDL has long resisted structural characterization at the molecular level, not only because of its size and complexity but because of the extensive heterogeneity in lipoprotein preparations [15–21]. The last decade has seen the emergence of cryo-electron microscopy (cryo-EM) as a viable approach for solving high-resolution membrane protein structures [22, 23]. In recent years, it has become possible to extend the molecular detail available from mid-resolution cryo-EM maps using the artificial intelligence encoded in programs like AlphaFold, which are often applied in tandem with structural refinement techniques involving molecular dynamics (MD) simulation [24–28]. Thus, with improvements in sample homogeneity and extensive computational sorting and classification of cryo-EM images, combined with advances in protein structure prediction and modeling, a detailed understanding of LDL molecular structure is now achieved.

## Results

### Cryo-EM reconstruction of human LDL

In preparation for structural characterization, LDL isolated from human serum by ultracentrifugation was purchased from a commercial vendor and further purified by size-exclusion chromatography (SEC). In addition to apoB100, gel electrophoresis revealed several smaller bands possibly corresponding to other exchangeable apolipoproteins as well as albumin, most of which were excluded by the SEC step (Extended Data Figure 1B-C). The slowest eluting fractions of LDL, corresponding to the smallest particles, were pooled, and prepared for cryo-EM imaging (Extended Data Figure 1A). We collected ∼4K micrographs and selected ∼600K LDL particles for further processing (Figure 1A-B, Extended Data Figure 2A-C, Extended Data Table 1). The particle diameters ranged from 16.2-22.4 nm (mean = 19.3 nm), suggesting the larger LDL species were successfully excluded, and their mean eccentricity was 0.48 (Figure 1C, Extended Data Figure 3A-C). Of these particles, ∼30% contained well-ordered stacks of high-density features in their core shown previously to correspond to the liquid crystalline phase of CE (Fig. 1B, Extended Data Figure 2,3) [19–21, 29]. These particles were excluded from the dataset as they biased the alignment unfavorably towards the core. A subset of particles that showed high-intensity features on their surface, interpreted as apoB100, were selected for further processing (Figure 1B, Extended Data Figure 2A). Extensive two-dimensional (2-D) and three-dimensional (3-D) classification resulted in a final reconstruction of the entire LDL particle at a global resolution of ∼9 Å from ∼53K images (Figure 1C, Extended Data Figure 2B, Extended Data Movie 1). At high contour levels, the map reveals a well-resolved (∼6-7 Å local resolution; Extended Data Figure 2D) globular protein domain connected to a continuous belt of high density that wraps around the particle’s circumference (Figure 1D). This belt divides the particle into two sides that when viewed from the “front” can be labeled the left (L) and right (R) faces (Figure 1D). At lower contours, additional features appear on the surface, until ultimately most of the exterior and interior are fully occupied. The large-scale morphology of the particle is semi-discoidal with polygonal features and dimensions of ∼17 nm x 20 nm (Figure 1E), placing it within the small dense LDL subclass [30]. The core is mostly disordered but retains a hint of liquid-crystalline CE packing, with the planar stacks oriented perpendicular to the curved face of the disc (Figure 1E). Reconstructions of different 3-D classes without any ordered CE were also obtained, yet no differences in peripheral features were observed (Extended Data Figure 2E). Lastly, using our full particle map, we generated a soft mask around the globular domain for focused refinement, achieving a reconstruction with local resolutions of up to ∼4.5 Å in this region (Extended Data Figure 2C).

**Figure 1.**
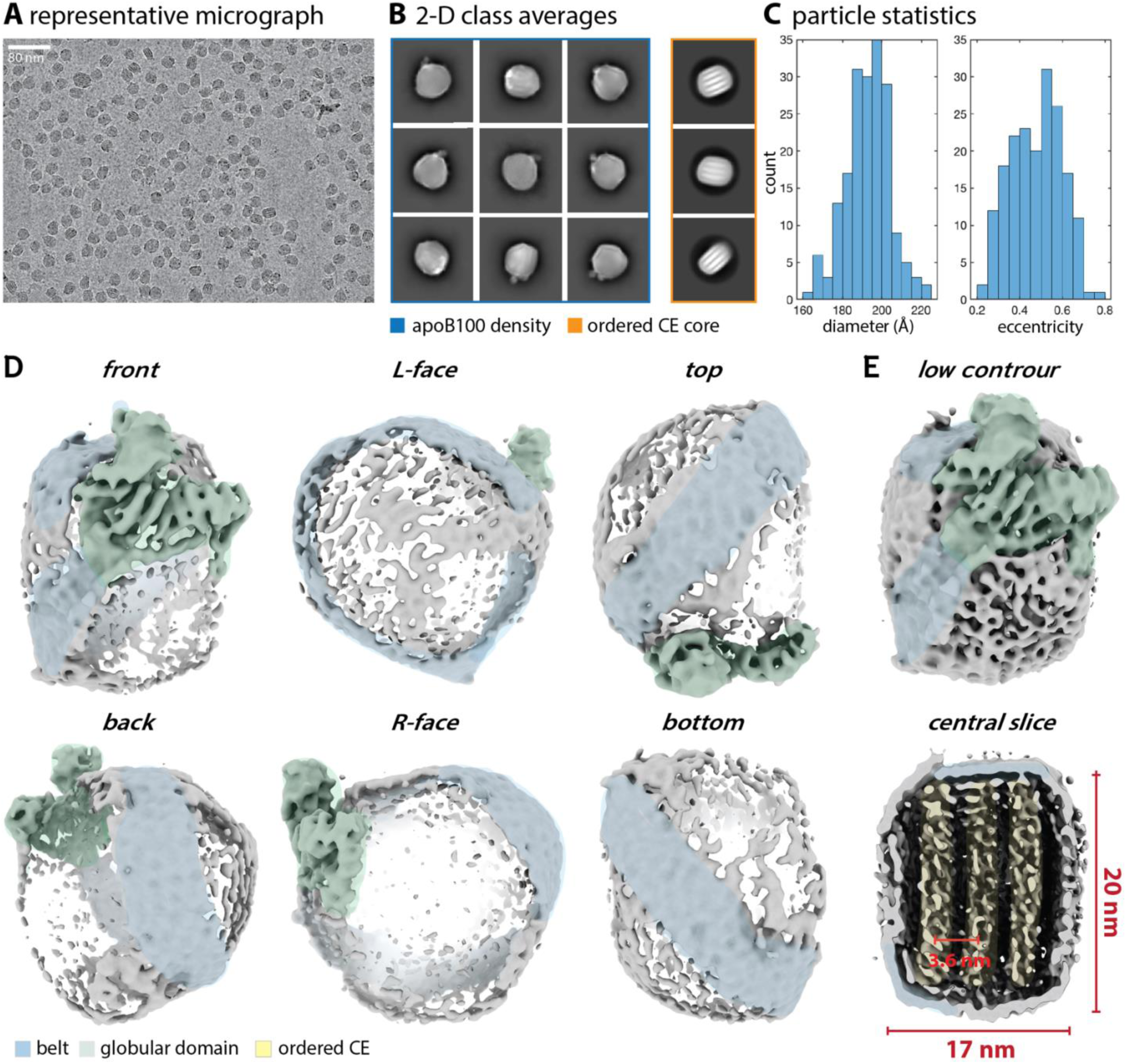
Cryo-EM reconstruction of human LDL. **A.** Representative cryo-EM micrograph showing vitrified LDL particles. **B.** Representative 2-D class averages showing a mixture of classes with clear apoB100 density around the particle exterior (blue) and particles with ordered CE cores (orange). **C.** Histogram of LDL particle diameters (average of the two principal axes) and eccentricities for 200 2-D class averages (see Extended Data Figure 2). **D.** 3-D reconstruction of the LDL particle at high contour with the globular domain and belt domain highlighted in green and blue, respectively, viewed from six different directions. **E.** Front view and cross-section through a low contour isosurface with particle dimensions and spacing between CE planes are indicated.

### Atomic model of apoB100

We next endeavored to construct an all-atom model of full-length apoB100 guided by our cryo-EM data. Given the lack of any existing high-resolution structural data, AlphaFold2 (AF2) [24] was employed to predict the complete 4563 residue structure of apoB100 as three contiguous overlapping fragments (Extended Data Figure 4A-C), which each exhibited reasonably high pLDDT confidence scores overall (Extended Data Figure 4A, D). When combined, the resulting structure could be divided grossly into two sections: (1) a globular N-terminal domain (NTD) ∼1000 residues in size and (2) a large C-terminal domain (CTD) consisting of a continuous ∼58-nm long x 4 nm wide β-sheet, which we term the β-belt, interspersed with 9 inter-strand inserts ranging in length from ∼30-700 residues (Figure 2A-B). The overall topology of the NTD and β-belt structures clearly matched the well-resolved regions of density in the cryo-EM map (Figure 1D, Extended Data Figure 5A). However, in the absence of the lipid particle, which could not be modeled with AF2, the predicted fragment models are collapsed into compact structures inconsistent with our data (Extended Data Figure 4A-C). We, therefore, used molecular-dynamics flexible fitting (MDFF) [27, 28] to refine the fragment conformations sequentially, starting with the NTD and then extending the model into the β-belt density (Extended Data Movie 2). These targeted regions matched our data well as evidenced by a high corresponding map-to-model cross-correlation (Extended Data Figure 5A) and provided a solid basis for interpreting the 9 more sparsely resolved inserts. Of these, the small inserts, 3, 5, and 7 (∼30-40 residues in length), which are predicted to be mostly unstructured, required little or no modification to fit the map, while the intermediate-sized inserts, 1, 2, 6, and 8 (∼90-170 residues in length), primarily required the use of MDFF to ‘flatten’ them out onto the surface. The two largest inserts, 4 and 9, which are 695 and 493 residues long, respectively, are both predicted to form strings of amphipathic helices that fold back on themselves into pairs separated by flexible loops, akin to the helical apolipoproteins like apolipoprotein A-1 (apoA-1) [31–33]. Several prominent regions of density in the cryo-EM map suggest that these inserts mediate long-range interactions across the LDL particle (Figure 1D) as described in greater detail below. We therefore employed a combination of MDFF and distance-based biasing potentials to systematically attract segments of these inserts with distal regions of the apoB100 structure whilst preserving their overall topology. We found close agreement between our final model and a comprehensive list of chemical cross-links reported in a recently published study [34] (Supplemental Information, Extended Data Figure 6). Taken together, our cryo-EM data and all-atom model establish for the first time the quintessential structural characteristics of full-length apoB100 in its native lipoprotein environment.

**Figure 2.**
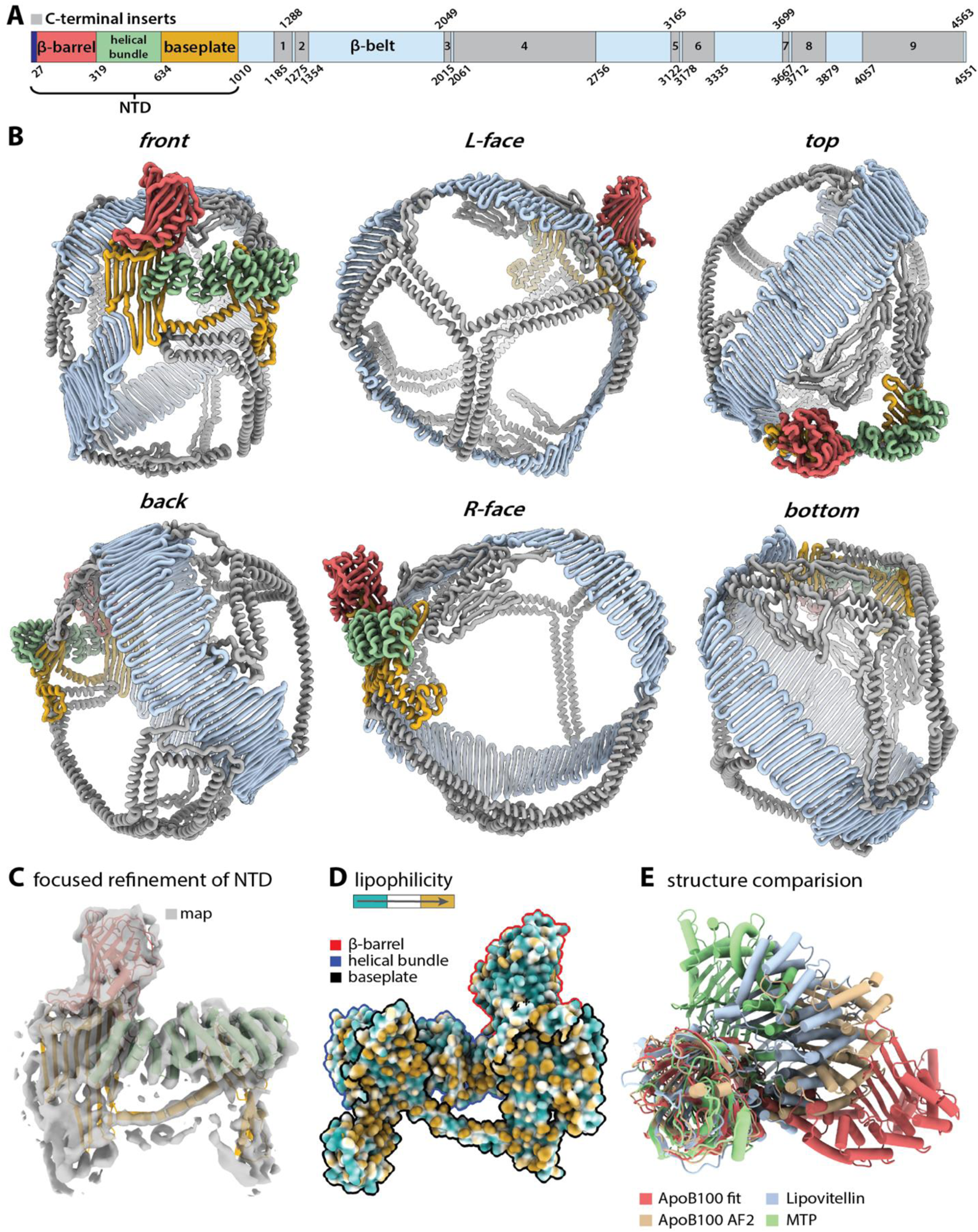
Atomic model of apoB100. **A.** Gene diagram of apoB100. **B.** Full atomic model of apoB100 viewed from six different directions and colored according to the gene diagram in panel A. **C.** Focused refinement of the NTD with relaxed atomic model viewed from the front. **D.** Atomic surface of the NTD viewed from the back and colored by lipophilicity with the baseplate outlined in black, the helical bundle in blue, and the β-barrel in red. **E.** Structural alignment of the relaxed NTD model with the unrelaxed AF2 model and the crystal structures of lipovitellin (PDB:1LSH) and MTP (PDB:6I7S).

### Structure of the apob100 NTD

The NTD can be divided into 3 subdomains that we term here: (1) the β-barrel (residues 28-319), (2) the helical bundle (residues 320-634), and (3) the baseplate (residues 635-1010) (Figure 2A-C). Residue 1010 was chosen to mark the end of the NTD because it corresponds to a small loop that separates the baseplate from the C-terminal β-belt. The first 27 residues correspond to a signal peptide, which is cleaved from the mature protein [6]. It should be noted that the residue numbering we use here includes the signal peptide. The β-barrel projects away from the surface of the particle at ∼35° and is the most prominent protein feature in the cryo-EM images (Figure 1B, Extended Data Figure 2A). It is composed of 11 strands and 3 helices, with one plugging the core of the barrel and another contacting the lipid surface, and it contains the smallest lipid-bound area of any domain (Figure 2D). The helical bundle is composed of 17 short helices and its arrangement can be described as a super-helical right-handed coiled-coil with a two-helix repeating unit [35] (Figure 2B-C). It lies partially on top of the baseplate with the central section contacting the lipid surface via 3 amphipathic helices (Figure 2D). The baseplate lies underneath the β-barrel and helical bundle and is composed of two amphipathic β-sheets separated by an amphipathic helix (Figure 2D). The first β-sheet is 6 strands long and has a 75 residue inter-strand insert between strands 4 and 5 composed of 3 amphipathic helices that project down from the baseplate. The second β-sheet is 8 strands long and roughly twice as wide as the first. The first 5 strands are the longest and extend upwards into the bottom of the β-barrel, while the last 3 are shorter and run in the opposite direction within the sheet. The NTD contains 7 of the 8 disulfide bonds in apoB100: 4 in the β-barrel, 2 in the helical bundle, and 1 in the baseplate (Extended Data Figure 7A).

### The NTD Is homologous to other LLTP superfamily members but adopts a more planar conformation on the surface of the particle

ApoB100 belongs to the large lipid transfer protein (LLTP) superfamily along with vitellogenin/lipovitellin, the main apolipoprotein of egg yolk, and microsomal triglyceride transfer protein (MTP), among others [36, 37]. MTP, which is essential for VLDL formation in the liver, is known to associate with apoB100 and is responsible for transferring lipids to the apoB100-containing nascent lipoprotein particle [38–41]. Both lipovitellin and MTP share ∼30-40% sequence similarity with the NTD and both have been previously crystallized, displaying similar conformations [35, 42–44]. At the domain level, our structure largely confirms previous homology models of the NTD based on these crystal structures [34, 45], however, it exhibits a unique conformation not yet seen in this class of proteins. Comparison of the relaxed NTD structure with lipovitellin and MTP yielded root mean squared deviations (RMSD) of ∼35 Å, with the largest deviations occurring in the helical bundle and baseplate (Figure 2E, Extended Data Figure 7B). In both lipovitellin and MTP, these regions exhibit a much tighter radius of curvature, which creates a self-contained lipid-binding cavity critical to their function. In line with this observation, the initial AF2 prediction of the NTD also exhibited a tighter radius of curvature, with some variation in curvature among the top 5 ranking models, which relaxed into a more planar conformation during refinement (Figure 2E, Extended Data Figure 7B). This predicted flexibility is further illustrated when the structure is segmented by the AF2 pae matrix [46], which divides the NTD in two down the center of the helical bundle and baseplate (Extended Data Figure 7C). Closer inspection of 3-D classification results from our cryo-EM data revealed some flexion in the NTD about this axis (Extended Data Movie 3), albeit not as pronounced, while anisotropic network modeling of the NTD produced a lowest-energy mode corresponding precisely to this same motion (Extended Data Figure 7D), suggesting it is intrinsic to the structure. Therefore, it seems plausible that the NTD of apoB100 could also adopt a more curved conformation with a lipid-binding cavity, for instance, during the early stages of protein synthesis and particle formation, as previously recognized [42, 45, 47, 48], but which can flatten out as the particle matures or curl back up as it is metabolized and shrinks.

### The highly conserved β-belt is the primary structural domain of LDL

The CTD of apoB100 can be subdivided into the β-belt (1774 residues) and 9 inter-strand inserts (1779 residues) (Figure 3A-B). The β-belt is the largest single domain in apoB100 and is highly conserved (Extended Data Figure 8A-B). It is composed of 120 strands between 3 and 16 residues long (mean = 11), with an average width of ∼4 nm (Figure 3B). This implies it cannot account for the full 17 nm width of the particle along the curved face (Figure 1D). Indeed, when viewed from the “front”, we see that the β-belt is tilted ∼35-45° relative to the long axis of the particle (Figure 2B). This tilt arises from the staggered arrangement of the first ∼20 β-strands which are followed immediately by the only sharp bend in the β-belt, a distinctive feature visible in the 2D class averages (Figure 1B, 2B, and Extended Data Figure 2A). This bend arises from the twisted arrangement of the β-belt in this region, which is present in the initial AF2 prediction and therefore not likely an induced conformation on the particle surface (Extended Data Figure 4A), implying it may have an important functional role, such as ensuring that the β-belt is properly oriented along the surface. After looping completely around the circumference, the β-belt passes back behind the full length of the NTD (Figure 2B). Although a few interactions are observed in our model between the NTD and β-belt, as well as between the NTD and inserts 6 and 8, these regions do not mesh tightly together and appear largely distinct within our cryo-EM map (Figure 2B). In light of this, we propose a mechanism whereby the β-belt can freely tighten and loosen, “sliding” past the NTD when needed, to accommodate different particle sizes. In addition to possessing this necessary flexibility, it has been shown that amphipathic β-sheets of apoB100 are critical structural domains that serve as sites for lipid incorporation during particle synthesis [49–51] and are strongly associated with the lipid membrane and/or hydrophobic core of particle [52–54]. This is consistent with the ∼55,720 Å^2^ of buried surface area observed in our structure. Along these lines, our cryo-EM data contains some evidence of coordinated lipids/CHS/CE/TG on the inner face of the β-belt at low-to-intermediate contour levels (Extended Data Figure 2E). Taken together, these results suggest that the β-belt is the primary structural domain of apoB100 responsible for maintaining LDL shape and cohesion.

**Figure 3.**
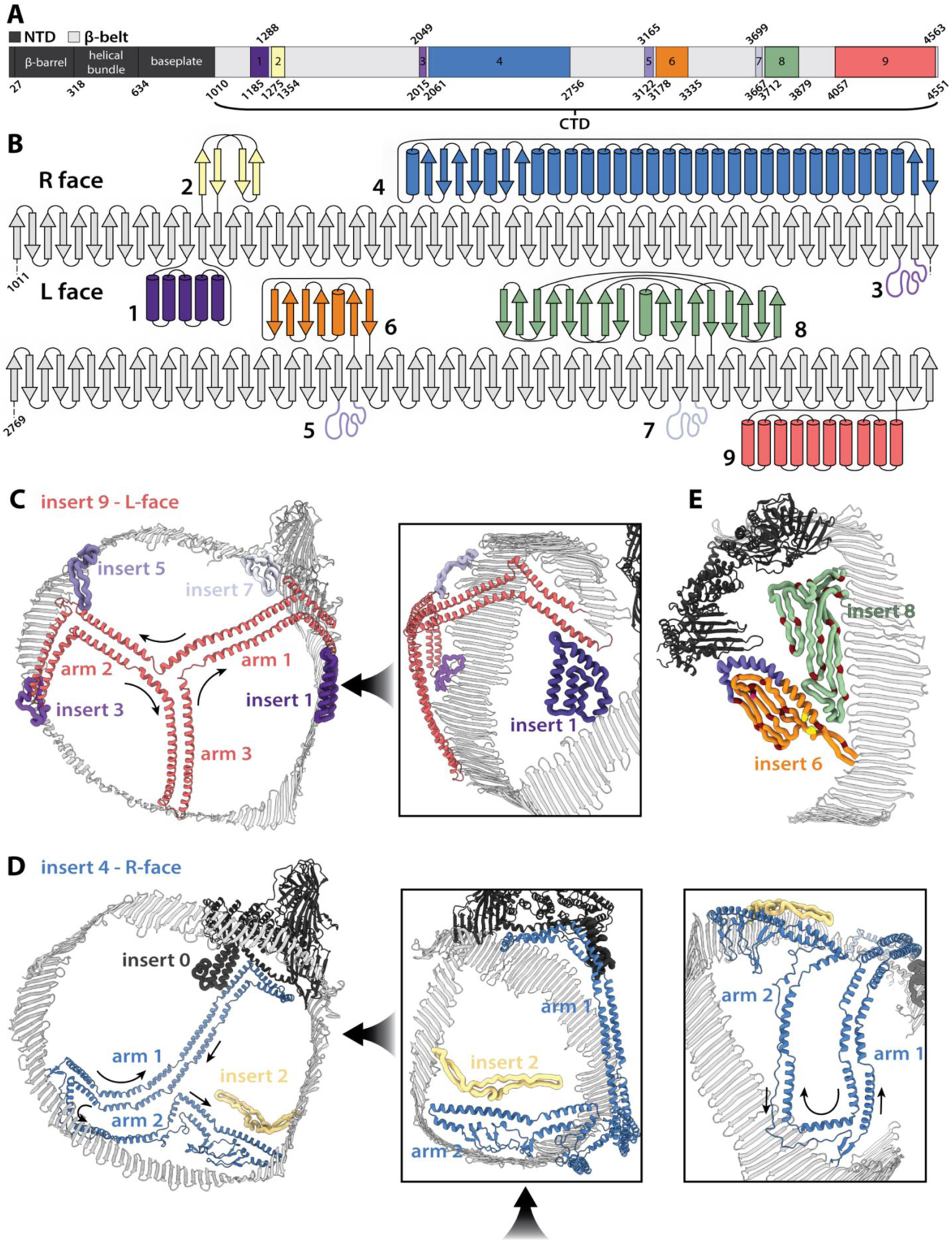
ApoB100 C-terminal inserts. **A.** Reproduction of the apoB100 gene diagram from Figure 1 with the 9 inserts colored uniquely, the NTD colored dark gray, and the β-belt light gray. **B.** Protein topology diagram of the C-terminal domain showing the continuous 120-strand β-belt and 9 inter-strand inserts with the R-face and L-face of the particle indicated. **C.** View of the L-face with insert 9 displayed as ribbons and inserts 1,3,5, and 7 as ropes. Black arrows point in the N-to-C direction of the protein chains **D.** View of the R-face with insert 4 depicted as ribbons along with insert 0 (baseplate insert) and insert 2 depicted as ropes. **E.** Inserts 6 and 8 are depicted as ropes with proline residues (maroon), fragment B3181[57] (purple), residue 3276 (pink), and disulfide bond (yellow) highlighted.

### The nine C-terminal inserts are arranged in evenly spaced pairs along the β-belt

Interspersed within the β-belt are the 9 inter-strand inserts, which are composed of ∼60% amphipathic α-helices, ∼30% coils, and ∼10% small amphipathic β-sheets (Figure 3B-C). They are on average less conserved than the other domains in apoB100 (Extended Data Figure 8). The smallest, inserts 3, 5, and 7, are mostly unstructured, while inserts 1 and 9 are predominantly helical, inserts 2 and 8 are predominantly β-sheets, and inserts 4 and 6 are a mixture of both. Five of the inserts project onto the L-face of the particle and four project onto the R-face (Figure 3B-C), and both faces contain one large helical insert along with several smaller ones. In addition, the first eight inserts are arranged along the β-belt in relatively evenly spaced pairs separated by a single β-strand and such that they project onto opposite faces (Figure 3B), suggesting a possible preference for symmetry across the two sides of the β-belt. However, despite this paired arrangement, there exists a relative imbalance across the two faces, in both the size and structure of the inserts. The L-face (687 residues) contains the 3 small unstructured inserts along with inserts 1 and 9, which are both entirely helical; the R-face (1081 residues) contains the three medium-sized inserts, 2, 6, and 8, which are predominantly β-sheets, and insert 4, which also contains a distinct β-sheet domain. This imbalance suggests the two faces of the particle may have unique properties and functional roles as described below.

### Insert 9 forms an expandable trefoil-shaped appendage and is the primary structural domain of the L-face

The L-face of the particle contains the small unstructured inserts 3, 5, and 7, the medium-sized helical insert 1, and the large helical insert 9. Insert 9 is located at the C-terminal tip of the β-belt, originating on the third to last β-strand on the L-face and returning on the last strand on the R-face, such that the β-belt is capped by two strands running in the opposite direction (Figure 3B). It is the only insert that does not have a pair on the opposite face. The L-face of our cryo-EM map shows a high-density feature loosely in the shape of a 3-point star with roughly linear segments separated by 120° (Figure 1D, Extended Data Figure 5B). Although AF2 predicted insert 9 to be a series of helical bundles that were folded back on themselves (Extended Data Figure 4A), the location of several small breaks suggested the existence of three distinct arms composed of pairs of helices (Extended Data Figure 9A). Indeed, once flattened out and relaxed into the map via MDFF, insert 9 was clearly arranged into a trefoil matching the distinct density feature just mentioned (Figure 3B, Extended Data Figure 5B), with the three arms meeting roughly in the center, creating the domed shape of the L-face (Figure 3C). Interestingly, when bound to the spherical form of high-density lipoprotein, apoA-1 also forms a trefoil [55]. If we label each arm by the order it appears in the sequence, we see that arms 1 and 2 are the longest (∼15-16 nm) and are composed of two pairs of helices separated by linkers that are bent at ∼50° and ∼90°, respectively, while arm 3 is the shortest (∼10 nm) and is composed of a single pair of helices. The arms of insert 9 are arranged perfectly to make long-range contacts at five roughly evenly spaced locations along the β-belt. The tip of arm 3 is positioned to contact the edge of the β-belt near residue 1720, while arm 2 is positioned to contact both inserts 3 and 5 at the tip and linker, respectively. The linker of arm 1 is positioned to contact insert 7 and insert 1 is positioned to contact the proximal segment of arm 1. This model for insert 9 is remarkably well-supported by chemical crosslinking data (Extended Data Figure 6A). When the L-face is viewed as a whole, it appears that insert 9 forms an expandable scaffolding-like appendage that reaches across the entire L-face and that inserts 1, 3, 5, and 7 are all positioned strategically along the β-belt to support it. The orientation of the proximal segment of arm 1 and its interaction with the β-belt and insert 7 are both predicted by AF2 with relatively high pae scores at the interfaces (Extended Data Figure 4D, 9B), suggesting the conformation observed here may represent the lowest energy state of apoB100.

### Insert 4 forms an extended prong-shaped appendage with two highly articulated arms and is the primary structural domain of the R-face

The opposing R-face of the particle contains inserts 2, 4, 6, and 8. Insert 4, which is the largest, forms an extended prong-shaped appendage with two highly articulated arms composed mostly of paired amphipathic helices separated by linkers. It occupies a region that extends from just below the NTD down through to the bottom right edge of the curved face of the semi-discoidal particle (Figure 2B, 3D). Consistent with having a more flexible structure, it is less well-resolved on average than insert 9, and the cryo-EM density is suggestive of a mixture of conformations. Arm 1 (346 residues) is the most uniformly helical, containing ∼8 pairs of helices, and it extends >30nm across the particle surface to contact the helical insert extending from the bottom of the baseplate (termed insert 0 to distinguish it from the β-belt inserts), as well as the helix separating the two β-sheets (Figure 3D). Instead of extending straight across the center of the R-face, it follows a curved path running roughly parallel to the β-belt, forming the bottom right edge of the particle (Figure 3D). Arm 1, the NTD, and the inner edge of the upstream β-belt enclose a circular region ∼14 nm wide that makes up the largest and flattest protein-free surface on the particle, which consequently has the weakest density in our cryo-EM map (Figure 1D). The base of the prong where the two arms meet sits adjacent to the β-belt, with the descending edge of arm 2 resting on top of it. This region is the best-resolved section of insert 4 and fits well into a prominent density feature in the map (Extended Data Figure 5C). The conformation of this segment is also present in the AF2 predictions, providing further support for our model (Extended Data Figure 4A, 9B), and again, suggesting this conformation may represent the lowest energy state. Arm 2 (327 residues) extends upwards between arm 1 and the β-belt where it can likely take on several conformations and interact with insert 2 (Figure 3D). Arm 2 is predicted to be less ordered and less uniformly helical. The ascending edge is composed of strings of helices like arm 1 and extends ∼24 nm across the particle surface, but at the distal end, a loosely ordered group of proline-rich β-hairpins flanked by short helices fold back onto the ascending edge, forming a somewhat cohesive domain. Although we could not unambiguously determine the orientation of this domain, its center-of-mass fit clearly into the prominent high-density feature at the “pointy” edge of the particle precisely where the β-belt makes a relatively sharp bend. Rigid docking of the isolated domain suggests two potential orientations, which differ by a ∼90° rotation along with an accompanying change in the positioning of insert 2, which, given its location, may either interact directly with arm2 or point back towards the NTD (Extended Data Figure 5C). It seems plausible that inserts 0 and 2 play a support role for insert 4 like the small inserts on the L-face, helping to anchor the two arms in place and/or coordinate their dynamics. Despite the reduced local resolution in this region of the map, our proposed structure is well-supported by the published chemical crosslinking data [34](Extended Data Figure 6). This data also shows evidence for a possible alternative conformation where arm 1 contacts insert 6 instead of the NTD. Although such a conformation would be physically possible, it is not supported by our cryo-EM data (Extended Data Figure 6B).

### Inserts 6 and 8 have similar structures and fall within the putative LDL-r binding region

The last two inserts, inserts 6 (156 residues) and 8 (166 residues), primarily occupy a region near the NTD in our model, forming several loose interactions with each other and the NTD. Overall, they are similar in size and structure, composed primarily of loosely folded amphipathic β-hairpins enriched in proline residues (Figure 3E). Insert 6 contains a single 35-residue-long α-helix in addition to the only disulfide bond outside the NTD (C3194:C3324), which links the ascending and descending strands near the base of the domain. Insert 8 has a small α-helix at the tip of the final hairpin, which contacts the helix of insert 6 in our model. Between the two, insert 8 is better resolved in our cryo-EM map, and fits clearly into a high-density feature positioned directly behind the NTD (Extended Data Figure 5C). This is further evident by the higher average pLDDT scores and the more uniform conformations among the five top-ranking AF2 predictions (Extended Data Figures 4A, 9C). The structure of insert 8 roughly matches the shape of the top boundary of the NTD, suggesting it may serve as a buffer between the NTD and β-belt, although its function is still unknown. Insert 6 on the other hand, is the most poorly resolved insert in our cryo-EM map (Extended Data Figures 2D, 5C), implying it is either dynamic, poorly folded, and/or loosely associated with the lipid surface. Along these lines, this region exhibited lower (∼40-50) pLDDT scores as well as alternative predicted conformations (Extended Data Figure 4A, C).

Although the function of insert 6, as with insert 8, is not apparent from the structure alone, current evidence suggests that the most probable location of the LDL-receptor (LDL-r) binding site (RBS) falls within the region loosely spanning these inserts, with some of the strongest evidence pointing to regions within insert 6 [5, 56–65]. For example, of the two most widely discussed proposed locations, termed Site A and Site B, Site A is positioned partially within insert 6 (residues 3173–3185), while Site B falls within the β-belt between inserts 6 and 8 (residues 3379–3395). A more recent study identified the fragment between residues 3209-3242 within the α-helix of insert 6 (Figure 3E) as the principal candidate for the RBS [57], while another study reported that an antibody whose epitope contains residue 3276, which is within the β-domain of insert 6 (Figure 3E), completely blocks LDL-r binding [56]. As a preliminary experiment, we used AF2 to predict the potential complex of LDL-r and insert 6, resulting in a model that located the binding site fully within the fragment spanning residues 3209-3242, albeit with only modest pae scores at the interface (Extended Data Figure 10A-C). It is believed that the RBS on apoB100 only becomes fully exposed after VLDL is converted to LDL and that this conversion is associated with a conformational change of apoB100 and/or the loss of other exchangeable apolipoproteins[64, 66–68]. It has also been reported that the small dense LDL subspecies, which are typically defined as particles with a diameter <22-25 nm, bind LDL-r with lower affinity [69–71]. This implies that the RBS may be presented sub-optimally in our structure, which has a diameter of only ∼20 nm. Given the positioning of insert 6 near insert 8 and the NTD, we expect that its local environment would change as the particle grows and shrinks in a way that could affect potential receptor binding, while the disulfide bond at the base would keep it properly folded. It should be noted that regions roughly corresponding to inserts 6 and 8 were previously identified based on their high proline content and susceptibility to protease digestion [54, 72]. This latter fact led the authors to conclude these regions may form compact structures that are unattached to the particle surface. This is consistent with our cryo-EM data in the case of insert 6, given its particularly poor resolution (Extended Data Figure 2D, 5D). Insert 8, on the other hand, is quite well-resolved and clearly bound to the surface; however, it is possible that when not constrained by the NTD on larger particles, it could diffuse more freely.

### A model for apoB100 conformational change

ApoB100 containing lipoproteins vary substantially in size, shape, and chemical composition; from the largest VLDLs (∼70 nm diameter, ∼220 nm circumference) to the smallest LDL (∼17 nm diameter, ∼53 nm circumference)[1], necessitating that apoB100 be flexible and robust enough to accommodate this wide range of conditions. Our structure suggests a potential mechanism for how the various domains of apoB100 act together in coordination to maintain particle shape and cohesion at different particle sizes (Figure 4). We propose the following four structural changes are able to occur in unison: (1) the NTD can curl up or flatten out to match the curvature of the particle, (2) the β-belt can tighten and loosen around the circumference, (3) insert 9 on the L-face can expand or compress by bending arms 1 and 2 about their flexible linkers while maintaining at least 2 additional points of contact along the length of the β-belt, and (4) insert 4 on the R-face can compress inward or straighten out by bending about its numerous flexible linkers all while maintaining its long-range contacts and orientation along the bottom right edge of the particle. While our structure paints a clear picture of how the L-face inserts might work together, the conformations and apparent functions of the R-face inserts are less clear. For instance, it is unclear what role inserts 6 and 8 might play in this process. On a larger particle, they would be freed from behind the NTD and available to potentially interact, either with each other or with other inserts like insert 4, as suggested by the chemical crosslinking data [34] (Extended Data Figure 6A). On a smaller particle, they would be pulled further behind the NTD, becoming even more isolated from the rest of the R-face. However, as described above, such changes to inserts 6 and 8 may be linked to their potential role in receptor binding and have little to no influence on the structural transitions required to maintain particle cohesion. Finally, when modeling the possible conformation of apoB100 on even larger lipoproteins such as VLDL, it is clear apoB100 would have less of an impact on their structure and lose the ability to form long-range interactions between inserts (Figure 4). In support of this hypothesis, no long-range chemical cross-links were observed within apoB100 associated with VLDL [34].

**Figure 4.**
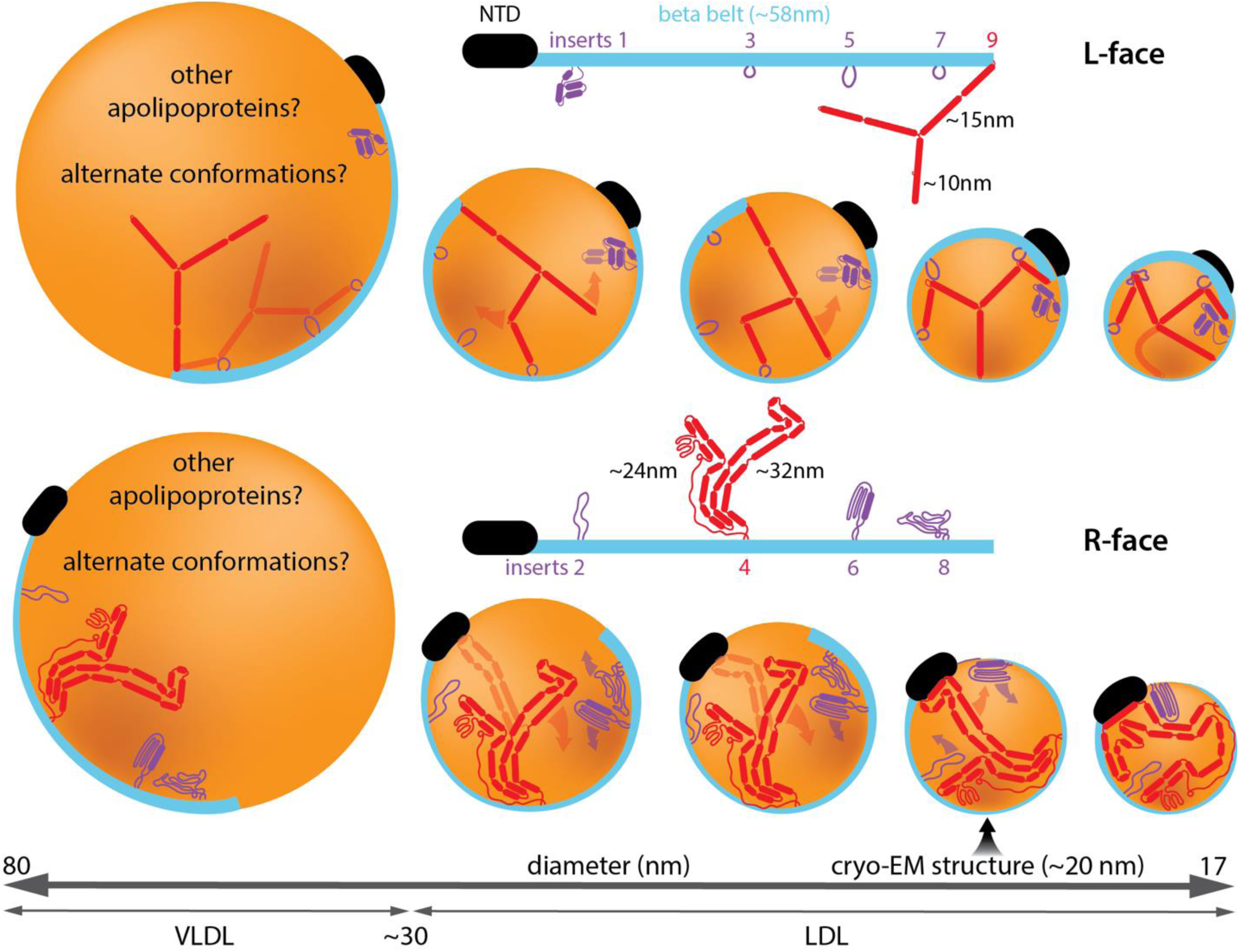
Model for apoB100 conformational change. A cartoon representation of different-sized apob100-containing lipoproteins, from VLDL to LDL, drawn roughly to scale depicting the proposed conformational changes that occur in apoB100 on the L-face (top) and R-face (bottom) of the particle. Transparent arrows and protein domains represent possible conformational changes and/or alternative conformations.

## Discussion

With the structure of apoB100 finally revealed, the past five decades of research can now be placed into the proper context. While such a monumental endeavor is beyond the scope of this study, we can address some of the important higher-level concepts. For instance, the prevailing structural model of apoB100, the pentapartite model, which has remained largely unchanged since its inception [54, 73], can now be updated along with the nomenclature. This model divided apoB100 into five domains termed βα1, β1, α2, β2, and α3. It was discovered early on that apoB100 contained primarily amphipathic secondary structural elements, and for the most part, the pentapartite model correctly distinguished between the β-sheet and α-helical content of the protein, although the exact boundaries were poorly defined [54]. βα1 corresponds to what had already been termed the NTD. Once the crystal structures of lipovitellin and MTP were solved, the exact boundaries of the apoB100 NTD were more clearly delineated based on homology modeling and differ only slightly from the ones we define here [54]. For the three sub-domains of the NTD, our nomenclature matches what has been used in the literature except for the third sub-domain, which we termed the baseplate, considering its role in supporting the other two sub-domains on the particle surface. Next, we found that β1 and β2, which were believed to form two independent domains, are part of the same continuous β-sheet, which we termed the β-belt. The idea of a belt-like structure has been proposed before [74, 75], though it was assumed the belt was broken by the large α2 domain. In addition, our structure reveals that α2 and α3 correspond to the large inter-strand inserts 4 and 9, respectively. Given their size and structural importance, it could be helpful if they retained unique domain names, in which case, we suggest simply renaming them α1 and α2 to better match the updated nomenclature. Considering the functions of inserts 6 and 8 are unknown, and inserts 1, 2, 3, 5, and 7 all appear to play more subtle roles in supporting the α domains, we suggest they be referred to simply by their insert numbering. If it becomes apparent that one or more of these inserts have a distinct functional role, for example, if insert 6 proves to be the receptor-binding domain, then the nomenclature can be updated.

The APOB gene encodes for both apoB100, and a truncated form called apoB48, a product of RNA editing that is composed of the N-terminal 48% of the protein only [76, 77]. ApoB48 is the primary apolipoprotein on chylomicrons, the largest class of lipoproteins (100s to >1000 nm in diameter) which are synthesized in the intestines and responsible for the packaging and transport of dietary-derived lipids[1]. From the structure apoB100, we can infer the structure of apoB48, which ends at residue 2179, placing the C-terminus within the fourth helix of insert 4, about halfway up the ascending edge of arm 1 (Extended Data Figure 11A). The possible function of this short segment of insert 4 is unclear, but perhaps it serves as a binding site for other exchangeable apolipoproteins. It is also not clear what functions inserts 1-3 would serve. It is clear, however, considering the size of chylomicrons, that apoB48 would have minimal impact on particle structure.

Metabolically, lipoprotein levels in the blood are tightly regulated, and when they become either too high or too low it can lead to serious illness [78]. In the context of atherosclerosis, the atherogenic potential of LDL is dose-dependent [9], therefore a significant medical emphasis is placed on maintaining levels as low as possible. In addition to being strongly influenced by diet and metabolic health [79], LDL levels are also affected by numerous genetic mutations [80]. For instance, there are >100 known disease-causing mutations in the APOB gene alone. Some of these cause elevated LDL levels, as seen in conditions like familial hypercholesterolemia (FH) and familial defective apolipoprotein B100 (FDB), while others may lead to lower levels, as in familial hypobetalipoproteinemia (FHBL)[12, 13, 81, 82]. In the case of FH, most mutations occur within the LDL-r gene, but there are several known FH mutations within apoB100, of which R3527Q is the most common and widely studied [82]. This is the defining mutation of FDB, which is a subtype of FH, and it is known to reduce the affinity of apoB100 for LDL-r. As such, it is widely believed that R3527 must either be within or near the RBS [54]. Our structure reveals that R3527 is located within the β-belt, about halfway between inserts 6 and 8, which we deduced was likely not within the RBS (Extended Data Figure 11B). In support of this, there is evidence showing that R3527Q does not directly block LDL-r binding but may cause a conformational change that disrupts the proper presentation of the RBS [61]. Considering our structure and the evidence placing the RBS within insert 6, the latter possibility seems more likely. Nevertheless, we cannot completely rule out the possibility that R3527 is within the RBS. For instance, the majority of other rare FH/FDB mutations also fall within this same stretch of β-belt, and most involve the loss of specific arginine residues, which are believed to be important for LDL-r binding [80] (Extended Data Figure 11B). Two other rare FH mutations map to insert 9, suggesting disruption of insert 9 structure or its long-range interactions could indirectly affect receptor binding (Extended Data Figure 11B). Mutations in APOB can also lead to dangerously low lipoprotein levels, like in the case of FHBL, which is most commonly caused by mutations in APOB resulting in C-terminal truncations of different sizes [83, 84]. Studies of truncated apoB100 mutants show that the length of the main β-sheet domains is directly related to lipoprotein particle size and indirectly related to particle density [51, 54]. This direct relationship to particle size is likely a result of insufficient lipid incorporation due to the truncated β-belt domain. The influence of the C-terminus (insert 9) is less clear, however. Although the particles appear normal, it has been shown that LDL containing apoB89, a common FHBL mutant resulting in the loss of insert 9 (Extended Data Figure 11B), are more rapidly cleared than wild-type particles, leading to the characteristic hypolipidemia. It was hypothesized that the loss of the C-terminus may lead to enhanced presentation of the LDL-r RBS [74, 85, 86]. Considering our structure, it is plausible that the loss of insert 9 could affect the conformation of the β-belt in a way that alters the presentation of the RBS. Perhaps when the long-range contacts mediated by insert 9 are abolished, the β-belt is no longer secured in its preferred orientation and becomes able to diffuse on the surface more freely, leading to increased exposure of the RBS while still maintaining particle cohesion.

The results presented here mark the beginning of a new era in LDL research where highly precise genetic, structural, functional, and computational studies are now possible. This newly acquired knowledge will not just advance our understanding of fundamental concepts related to lipid and cholesterol metabolism but will accelerate the design of potential next-generation LDL-modulating therapies for the treatment and prevention of atherosclerosis, such as monoclonal antibodies that can target LDL, or even specific subclasses of LDL, directly to achieve more precise treatment with potentially fewer side effects [87].

## Materials and Methods

### Sample Preparation

∼500 ug of human LDL purified by density gradient ultracentrifugation was purchased from Thermo Fisher (Catalog number: L3486) and then further purified via size exclusion chromatography using a Superose 6 Increase column (Cytiva) and Akta Pure system (Cytiva). To reduce heterogeneity in LDL size prior to cryo-EM, the fractions corresponding to the slowest eluting quarter of the LDL peak (rightmost tail) were pooled and concentrated to ∼1 mg/ml. 4 µl of this sample was then applied to plasma-cleaned holey carbon grids (Quantifoil) and plunge frozen with an FEI vitrobot mark IV at 4°C and 100% humidity.

### SDS-PAGE

Sodium dodecyl sulfate-polyacrylamide gel electrophoresis (SDS-PAGE) was performed in a Bio-Rad Mini-Protean Tetra Cell chamber using 4–20% gradient precast tris-glycine gels. ∼1µg of LDL was loaded per lane as measured by apoB100 absorbance at 280nm.

### Cryo-EM data collection

Imaging was performed on an FEI Titan Krios G4 Cryo-TEM (Thermo Fisher) equipped with a Gatan K3 direct detector and Gatan BioQuantum Imaging Filter. Automated data collection was controlled using SerialEM[88]. Detailed imaging statistics can be found in Extended Data Table 1.

### Cryo-EM data processing

All data processing was performed with CryoSparc v4.0[89] following the workflow presented in Extended Data Figure 2. Briefly, ∼600K LDL particles were picked and extracted after frame alignment, dose-weighting, CTF estimation, and micrograph curation. Numerous rounds of 2-D classification followed by subset selection were performed, selecting for homogeneous particles based on size and the presence of visible apoB100 density. Particles with ordered stacks of CE in their core were excluded from further processing. Numerous rounds of 3-D classification with and without masks around the NTD using different algorithms were performed followed by subset selection, resulting in a final set of ∼52K homogeneous particles that refined to a global resolution of ∼9 Å. All 3-D refinement jobs were performed with the non-uniform refinement algorithm. Lastly, a soft mask was generated around the NTD of apoB100 and used for local refinement, resulting in a ∼6 Å-resolution (global) reconstruction of the NTD. Numerous alternative processing pipelines were explored with only minor differences in the final appearance and resolution. Local resolution analysis was performed in CryoSparc. All map and model visualizations were generated with ChimeraX [46, 90].

### Morphological analysis

Morphological analysis of 2-D class averages was performed in MATLAB v9.12.0.2009381 (R2022a) equipped with the Image Processing Toolbox v11.5 (The MathWorks Inc). Class averages were generated from the particle stack derived from the first round of classification and subset selection, prior to excluding particles with ordered CE core density.

### Model Building and Refinement

All protein structure predictions presented in the paper were carried out with AlphaFold2[24] as implemented through ColabFold [91]. Due to current single-prediction size limitations, the apoB100 sequence (Uniprot P04114) was divided into three fragments (residues 1-1820, residues 1681-3500, and residues 3361-4563) with 140-residue overlaps to assist with subsequent model concatenation. The full-length apoB100 model was constructed progressively through a series of molecular dynamics (MD)-based structural refinement simulations, utilizing particularly the molecular dynamics flexible fitting (MDFF) methodology [28]. When modeling the insert structures, the collective variables (Colvars) module [92] was employed to create appropriate biasing potentials between distant regions of the apoB100 structure, which were used in tandem with MDFF. To refine model stereochemistry, the structure was first subjected to a series of restrained equilibration simulations and conjugate gradient energy minimizations in explicit solvent, followed by real-space refinement in Phenix v1.20 [93] and, finally, manual refinement using ISOLDE [94].

Model building and visualization were done using a combination of VMD v1.9.4 [95] and ChimeraX v1.6.1 [46, 96]. All simulations were performed using NAMD v3.0 [97, 98] and the CHARMM36 force field [99]. MDFF simulations were performed using standard settings [28] with a coupling factor of 0.3 applied to the protein backbone. To help overcome regions of heterogeneous resolution in the cryo-EM data, we employed a so-called cascade approach [27] in each MDFF simulation, first fitting the target region to four Gaussian-blurred maps (more blurred progressing to less blurred) before finally fitting to the original map. Harmonic restraints were applied during all fittings to maintain secondary structure and to prevent the formation of cis-peptide bonds and chirality errors.

### Normal Mode Analysis

The anisotropic network model of the apoB100 NTD (residues 28-1010) backbone atoms was created in ProDy [100, 101] and used to calculate the 10 lowest-energy normal modes of the structure. The results were visualized using the normal mode wizard [100] implemented through VMD [95].

## Data availability

Cryo-electron microscopy maps and refined atomic models will be deposited into the appropriate public databases upon publication of the peer-reviewed article. Extended data movies for this preprint can be found at: https://zenodo.org/doi/10.5281/zenodo.10723756

## Code availability

The Matlab code for performing morphological analysis of 2D class averages can be found at GitHub: https://github.com/ZTBioPhysics/morphological-analysis-of-LDL-cryo-EM-2D-class-averages.git

## Acknowledgments

Z.T.B. would like to thank the staff of the Mizzou Electron Microscopy Core, especially Dr. Min Su, and the Mizzou Research Support Solutions staff, especially Matt Stanley and Asif Ahamed Magdoom Ali, for their technical support with microscopy and high-performance computing, respectively. Computational aspects of this work were partially supported by the high-performance computing infrastructure provided by Research Computing Support Services and in part by the National Science Foundation under grant number CNS-1429294 at the University of Missouri, Columbia MO. Z.T.B would like to thank Anvitha Boosani for running SDS-PAGE gels. We would also like to thank Steven Van Doren, Michael Chapman, and Gavin King for critical reading of the manuscript.

## Author contributions

Z.T.B. conceived of the experiments, prepared samples, performed electron microscopy data collection and processing, model building, NMA, sequence analysis, interpreted results, prepared figures, and wrote the manuscript. C.K.C. performed model building and validation, molecular dynamics simulations, and flexible fitting simulations, interpreted the results, and wrote the manuscript.

## Competing interests

The authors declare no competing interests.

## Funding

This work was supported by University of Missouri faculty startup packages for Z.T.B. and C.K.C. from the departments of Biochemistry and Physics, respectively.

## Extended Data Figures

**Extended Data Table 1.**
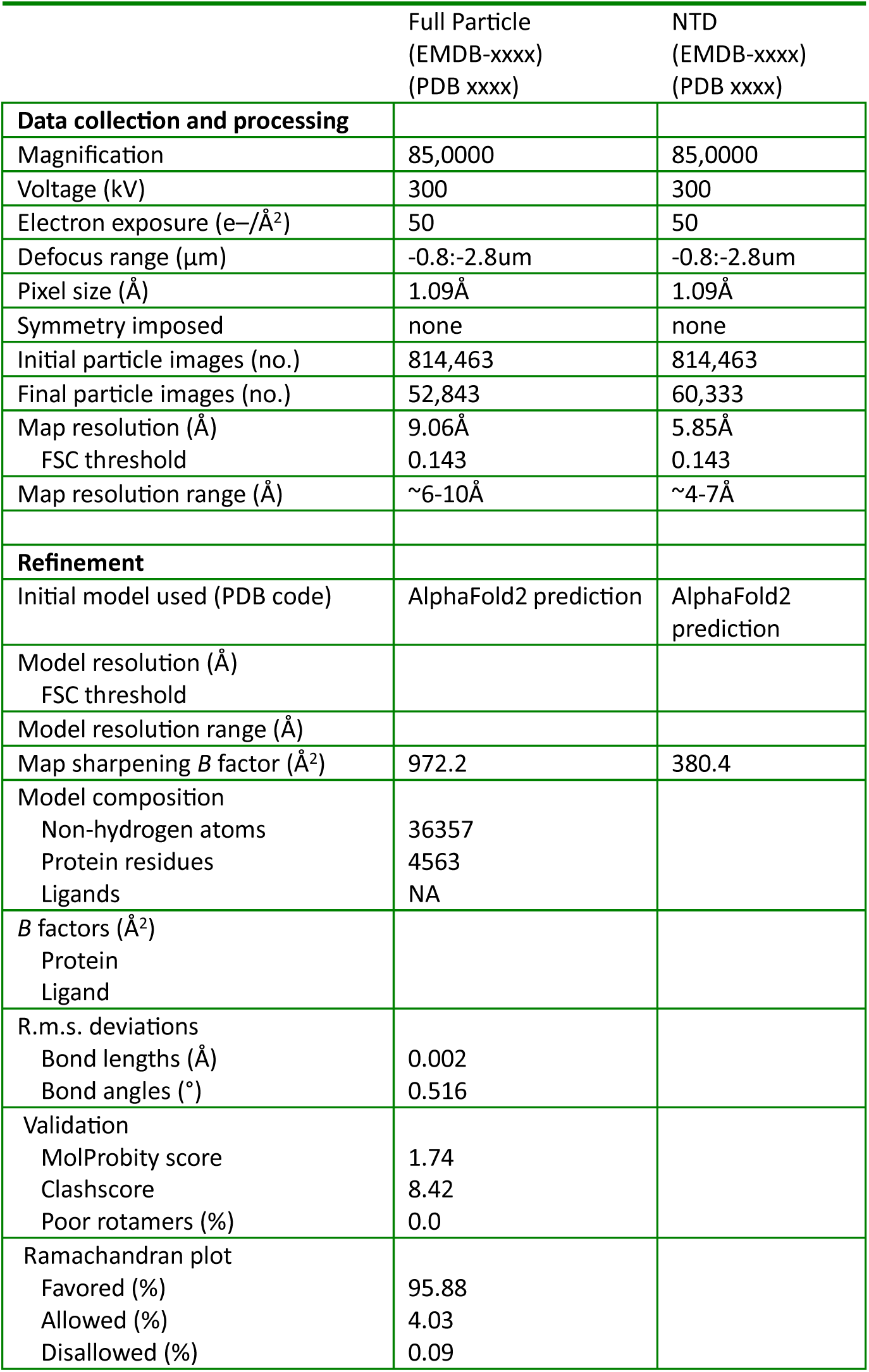
Cryo-EM data collection, refinement, and validation statistics.

**Extended Data Figure 1.**
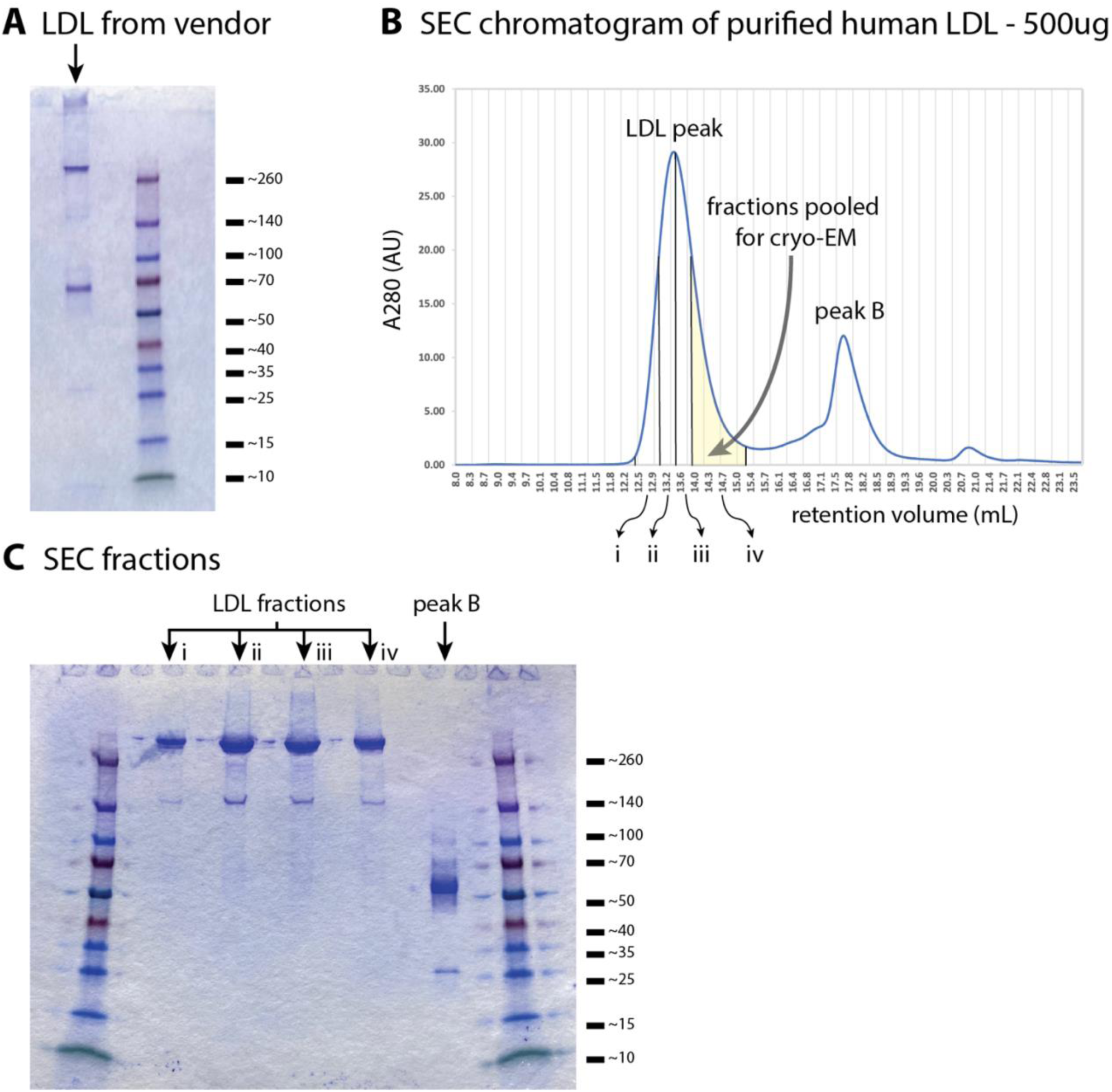
SDS-PAGE and SEC of purified LDL. **A.** Sodium dodecyl sulfate-polyacrylamide gel electrophoresis (SDS-PAGE) and (**B**) SEC chromatogram of human LDL purified by density gradient ultracentrifugation (purchased from commercial vendor). The LDL peak is indicated, and the fractions collected for cryo-EM imaging are highlighted in yellow. **B.** SDS-PAGE gel of isolated SEC fractions.

**Extended Data Figure 2.**
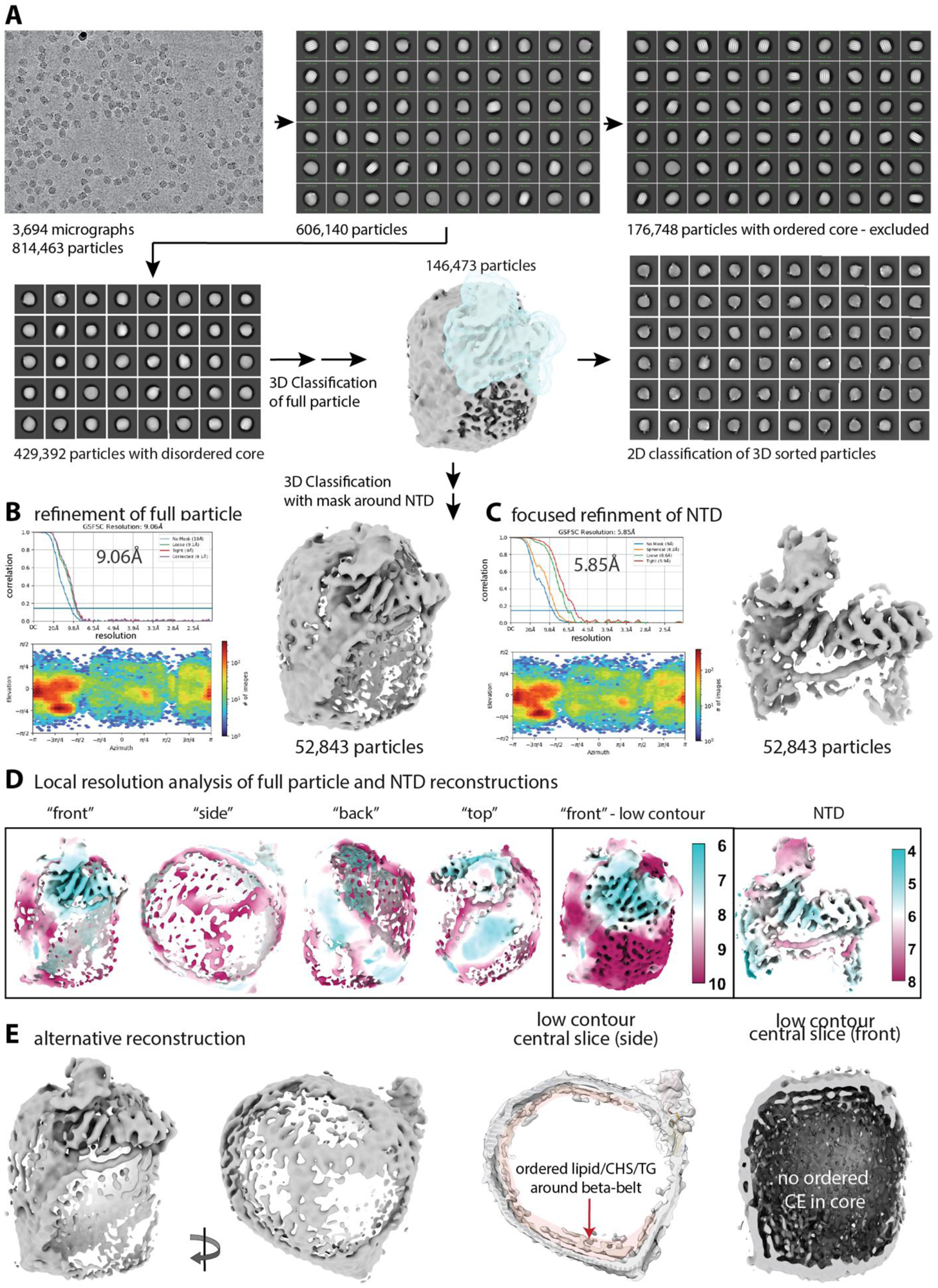
Cryo-EM data processing workflow and resolution estimates. **A.** Data processing workflow showing a representative micrograph, 2-D class averages, and intermediate 3-D reconstructions. **B-C.** Fourier shell correlation curves with global resolution estimate and angular distribution heatmaps for the full LDL particle and focused refinement of the NTD. **D.** Local resolution estimates for both maps. **E.** Alternative reconstruction of the full LDL particle that does not contain any liquid crystalline CE viewed from 2 directions and as central slices at low contour. Also shown is the density potentially corresponding to coordinated lipids/CE/CHS/TG on the inner face of the β-belt.

**Extended Data Figure 3.**
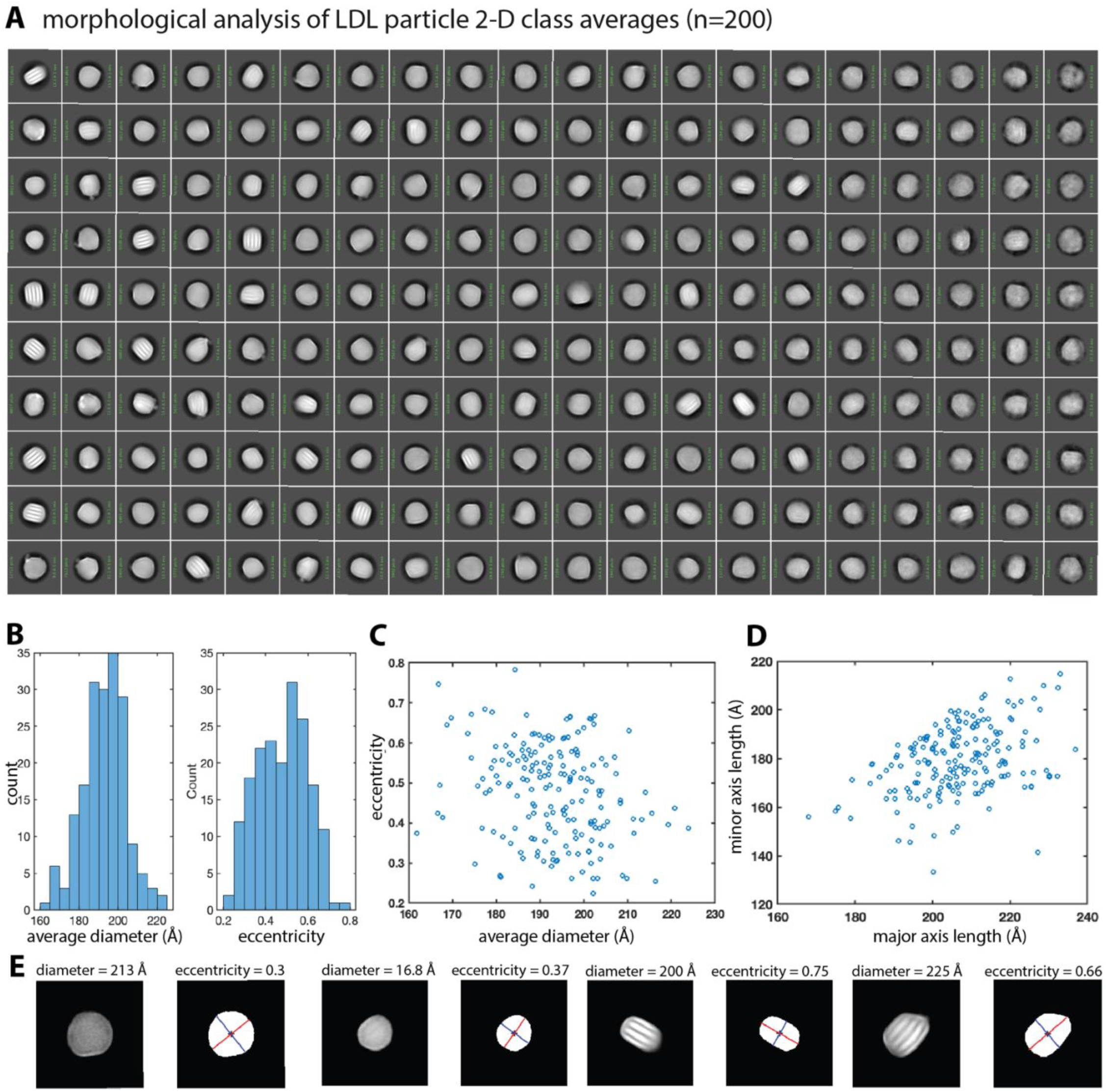
Morphological analysis of 2-D class averages. **A.** 200 2-D class averages of LDL particles used for morphological analysis. **B.** Histogram of average particle diameters and eccentricities. **C-D.** Scatter plot showing average particle diameter vs. mean eccentricity and major vs. minor axis length. **E.** Representative class averages from different extremes of the distribution.

**Extended Data Figure 4.**
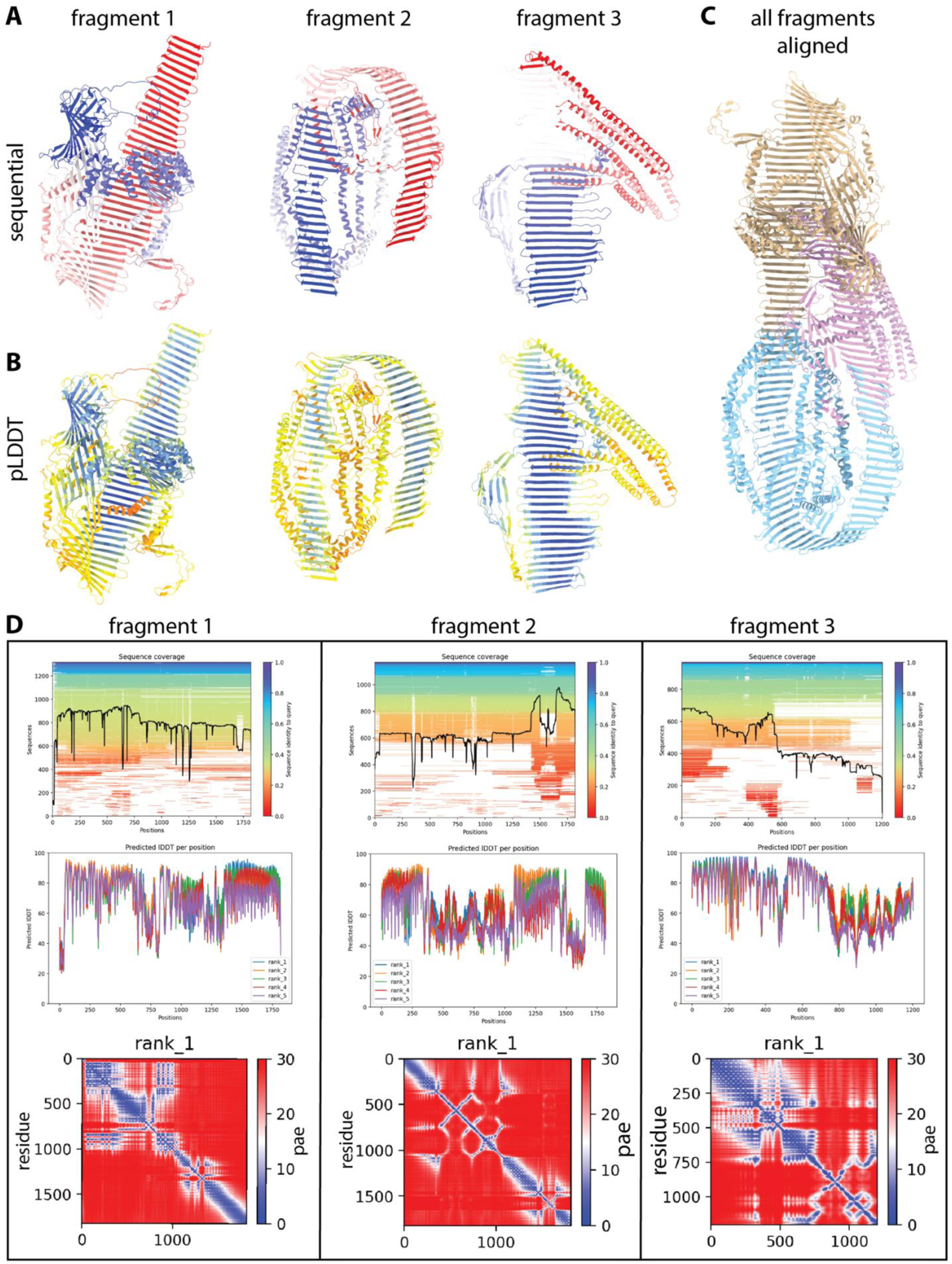
AlphaFold2 structure prediction for apoB100. **A.** AF2 predicted structures for the three contiguous apoB100 fragments colored sequentially and by (**B**) pLDDT score. **C.** All three AF2 fragments aligned to one another using overlapping sequence region. **D.** Per-residue sequence coverage, pLDDT score, and pae score for each fragment.

**Extended Data Figure 5.**
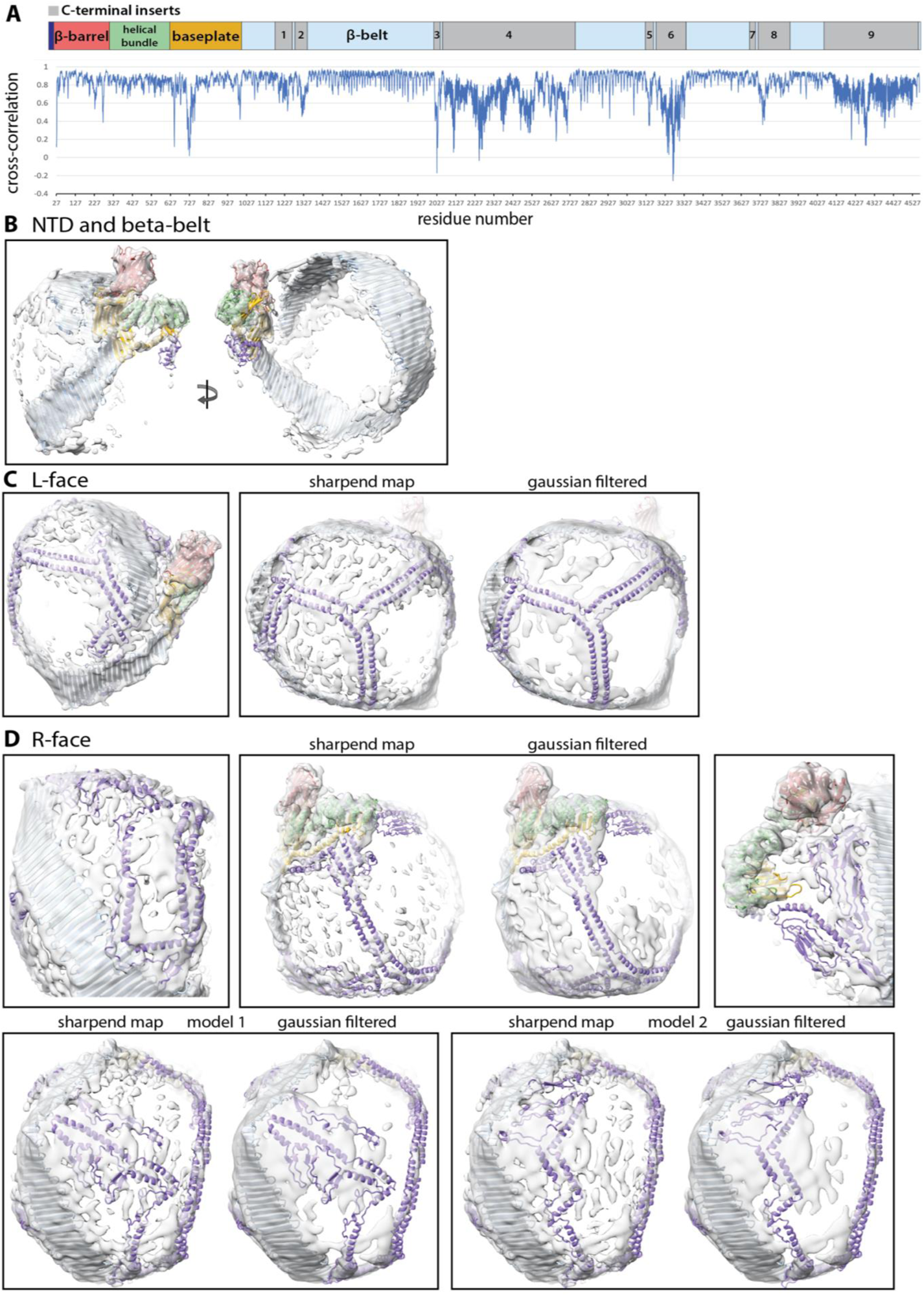
Map-to-model fits. **A.** Per-residue map-to-model cross-correlation plot. **B-D.** ApoB100 atomic models fit into the cryo-EM map containing just the NTD and β-belt at high contour, or the full protein viewed from the L-face and R-face at low contour. A Gaussian-filtered map is also shown for the L and R-faces.

**Extended Data Figure 6.**
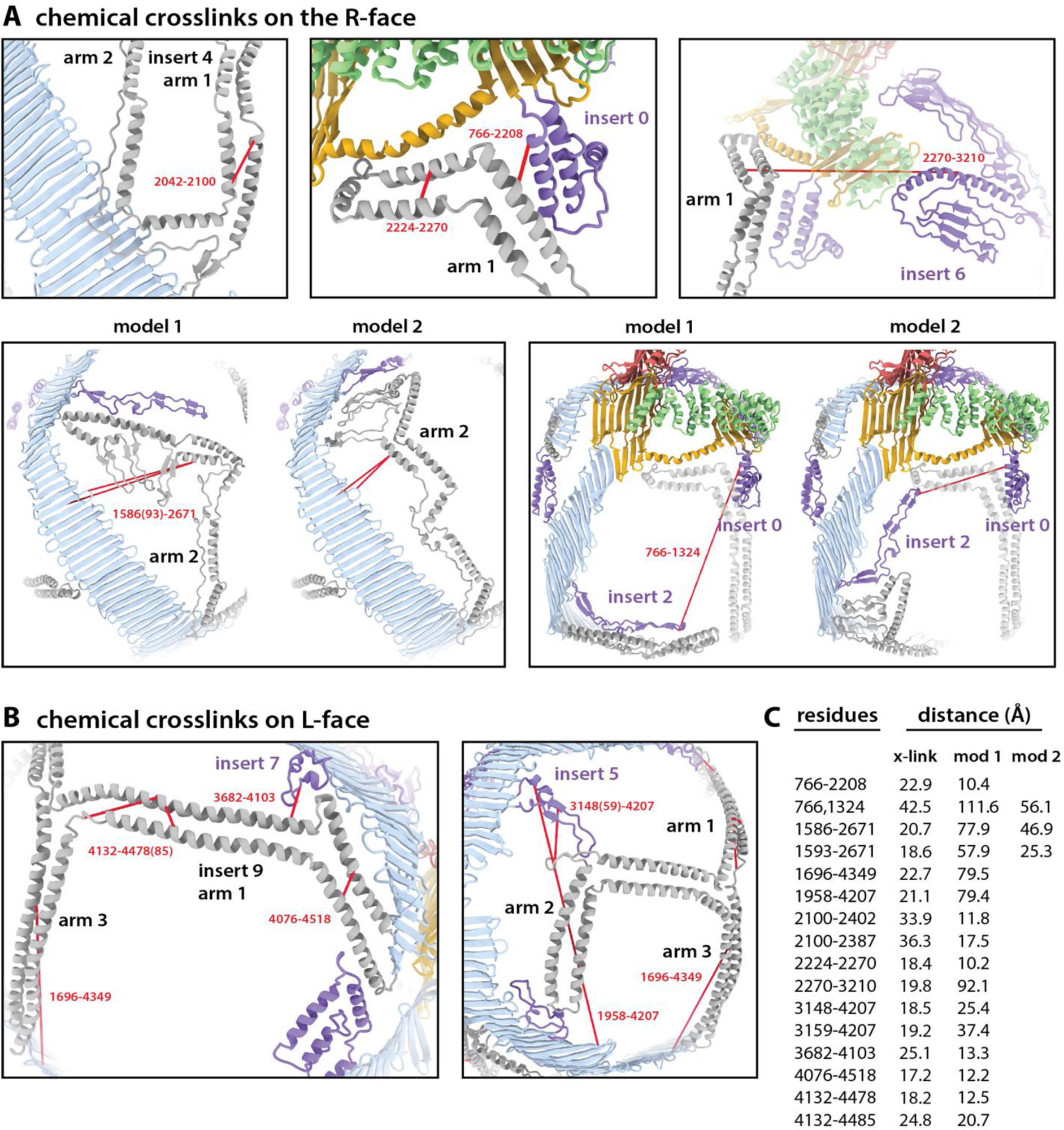
Long-range chemical cross-links mapped to the apoB100 structure. **A-B.** Experimentally determined long-range chemical cross-links within the C-terminal inserts mapped to the apoB100 structure on the R-face and L-face. Note that model 1 and model 2 differ only in their placement of arm 2 of insert 4. **C.** List of all chemical cross-links and their associated distances reproduced from the published study [34] and measured from the two atomic models.

**Extended Data Figure 7.**
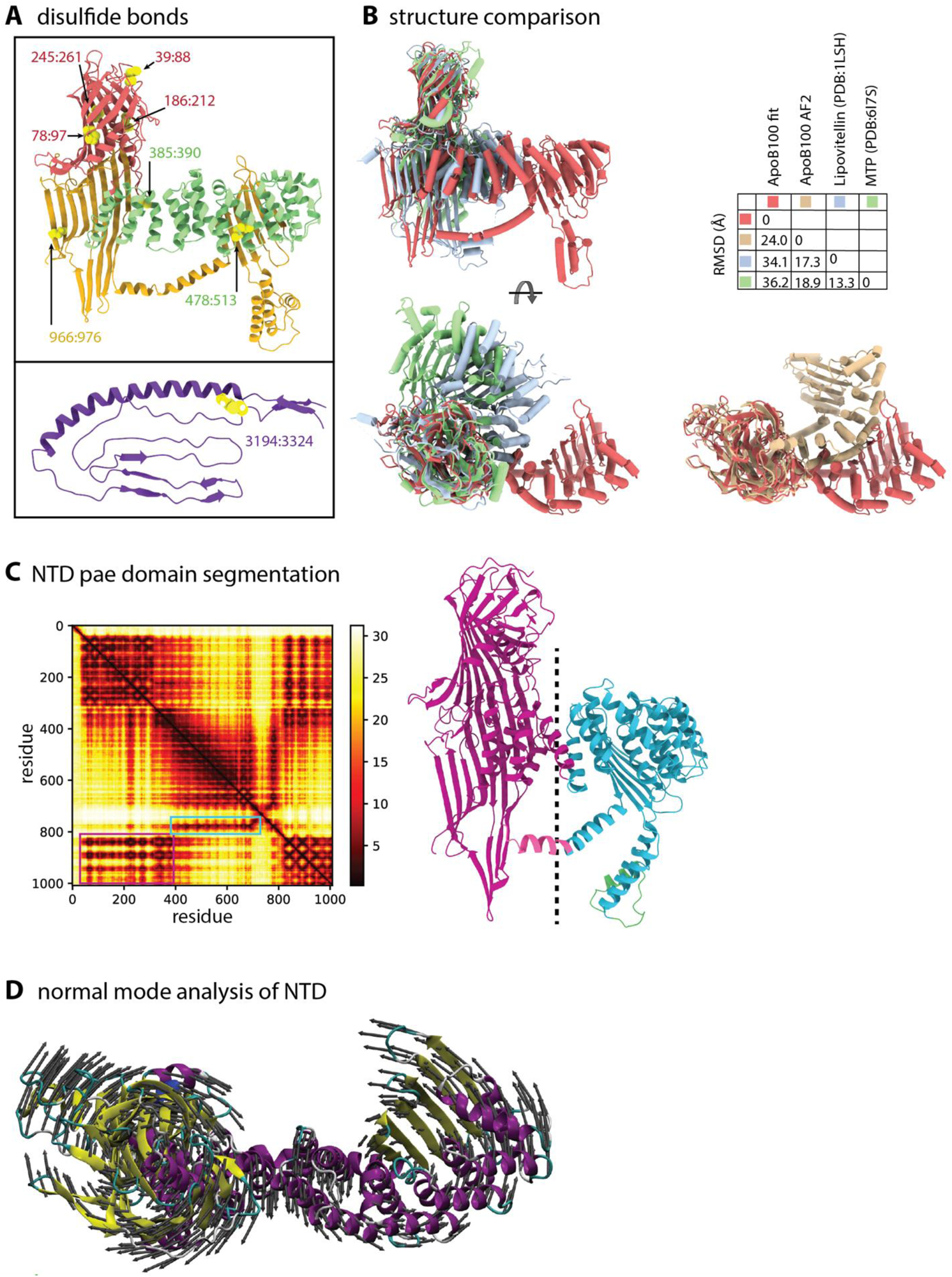
Extended structural analysis of apoB100 NTD. **A.** ApoB100 disulfide bonds in the NTD (top) and insert 6 (bottom). **B.** Expanded structural comparison between the predicted and relaxed models of the apoB100 NTD and the crystal structures of lipovitellin and MTP. **C.** AF2 pae matrix for the NTD alone and the automatic structural segmentation of the NTD derived from the pae matrix as implemented in ChimeraX. **D.** Top view of the NTD colored by secondary structural element showing the vectors of motion (gray arrows) calculated from the slowest normal mode.

**Extended Data Figure 8.**
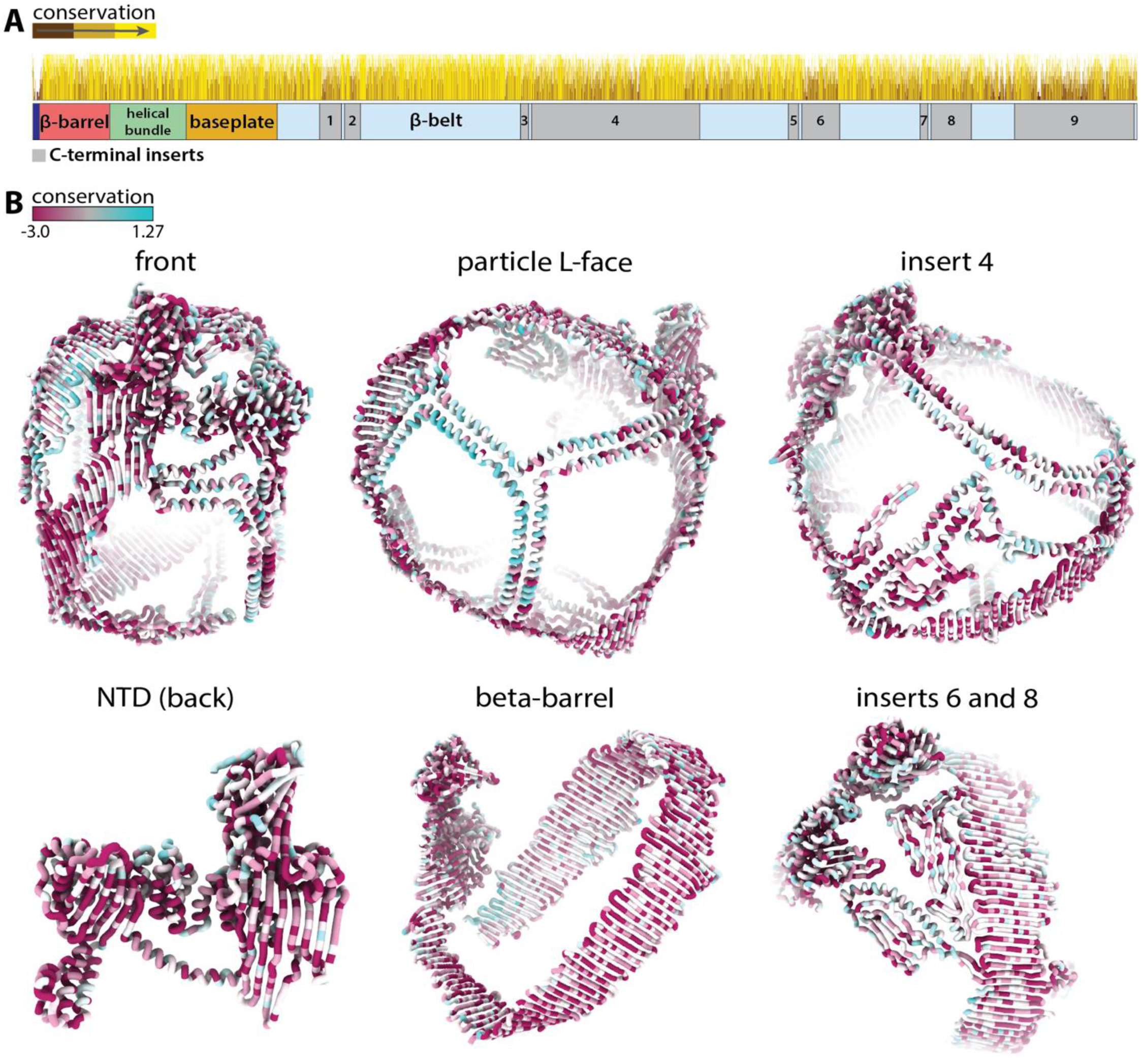
Sequence conservation of apoB100. **A.** Per-residue sequence conservation score calculated from a multiple-sequence alignment of apoB100 variants plotted on top of the apoB100 gene diagram. The alignment was calculated with UniProt BLAST [102] and visualized with Jalview [103]. **B.** Sequence conservation score mapped to the apoB100 atomic model using ChimeraX.

**Extended Data Figure 9.**
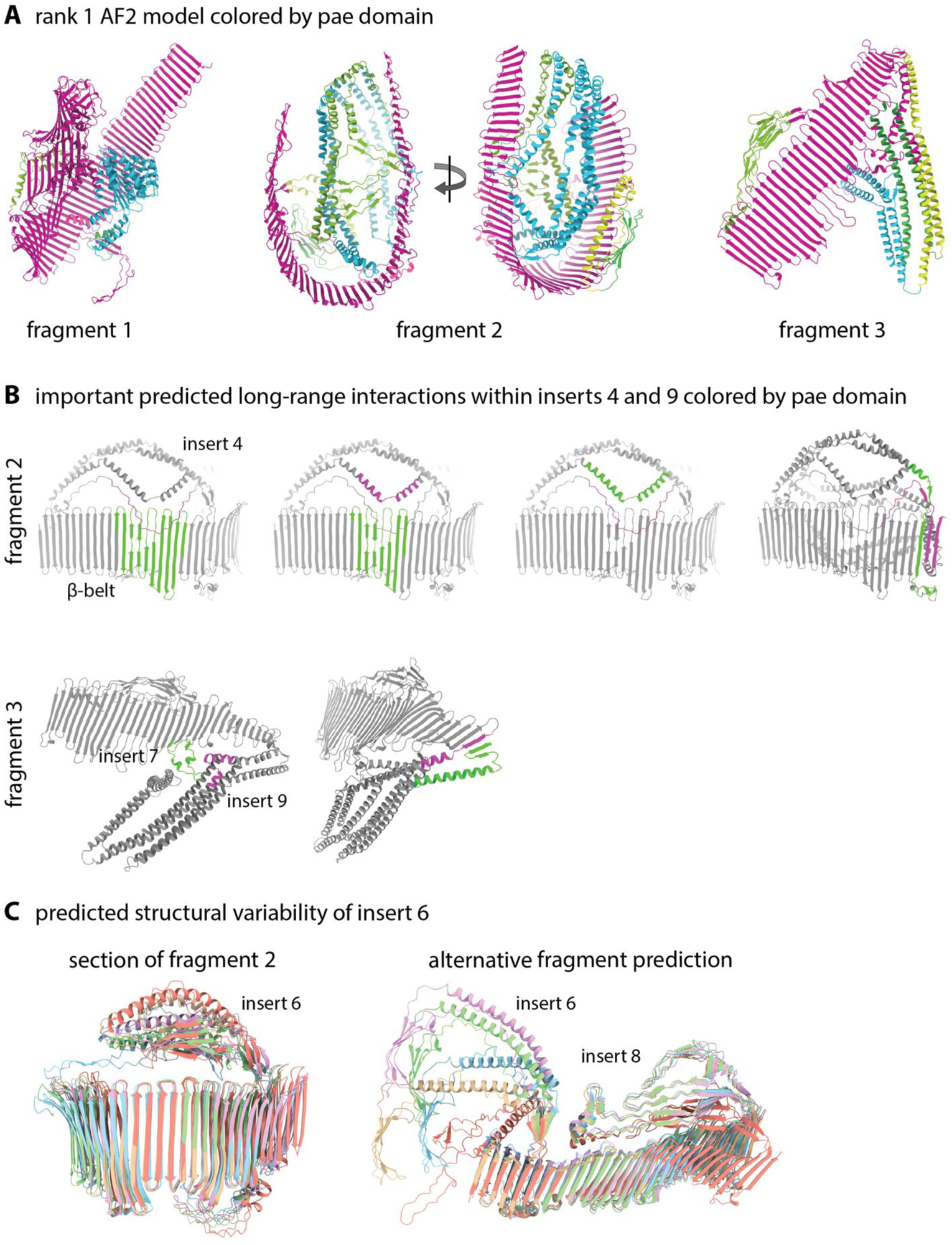
Extended AF2 structure analysis. **A.** The three predicted fragments of apoB100 colored by “pae domain” as implemented through ChimeraX. **B.** Important predicted long-range tertiary contacts in helical inserts 4 and 9. **C.** The five top-ranking AF2 models for the region containing insert 6 from fragment 2 along with the region containing inserts 6 and 8 from an alternative prediction.

**Extended Data Figure 10.**
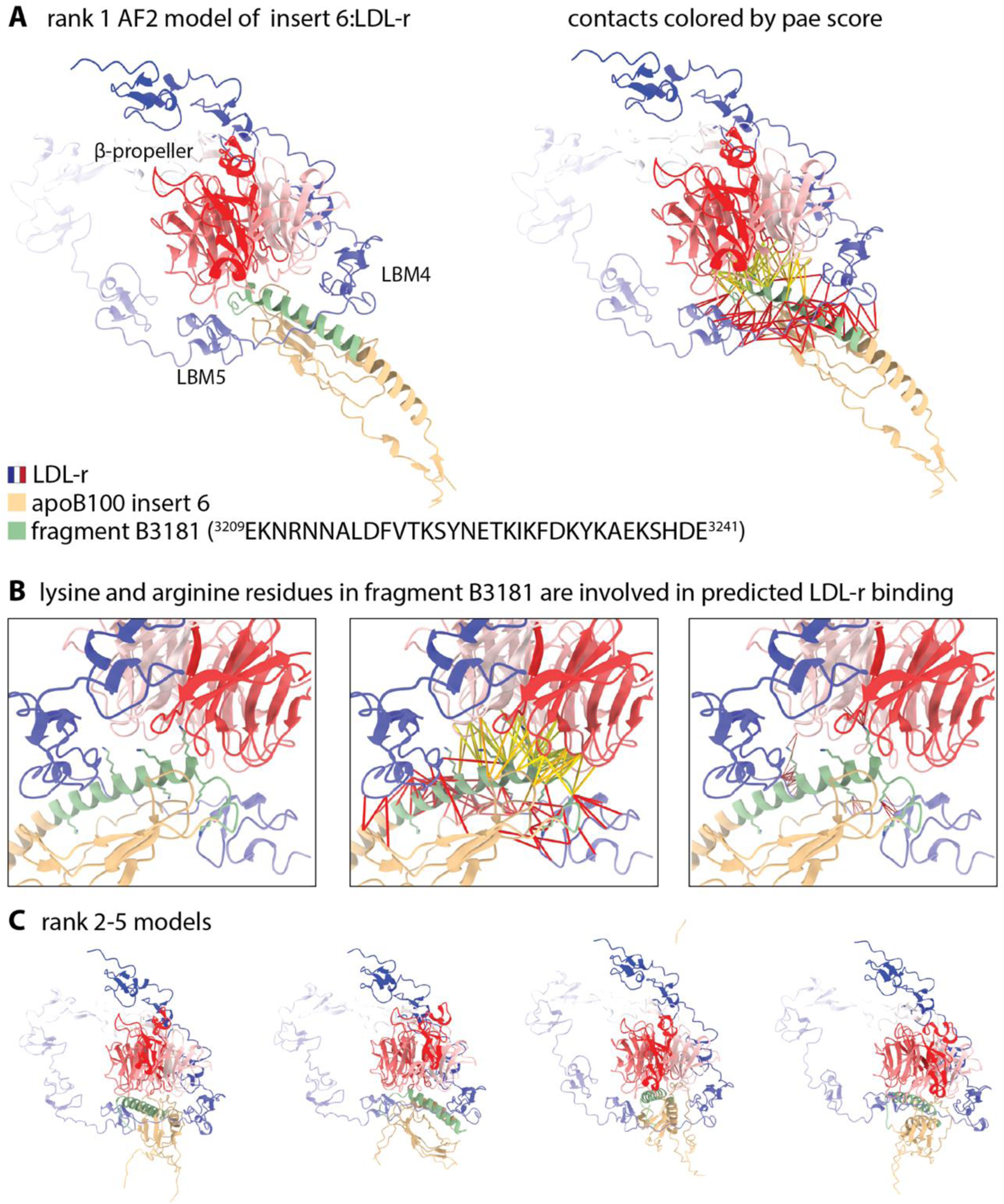
AF2 modeling of LDL-r in complex with insert 6. **A.** The top-ranking AF2 prediction of LDL-r (colored sequentially) in complex with insert 6 (tan). The residues corresponding to fragment B3181[57] are colored green. Also shown (right) are the predicted contacts (within 6 Å) between the two protein chains colored by relative pae score (yellow = lower pae and higher confidence). **B.** Close-up view of the predicted interface with lysine residues predicted to be within the LDL-r binding site shown with and without the contacts colored. **C.** The four other top-ranking predictions showing alternative conformations of insert 6.

**Extended Data Figure 11.**
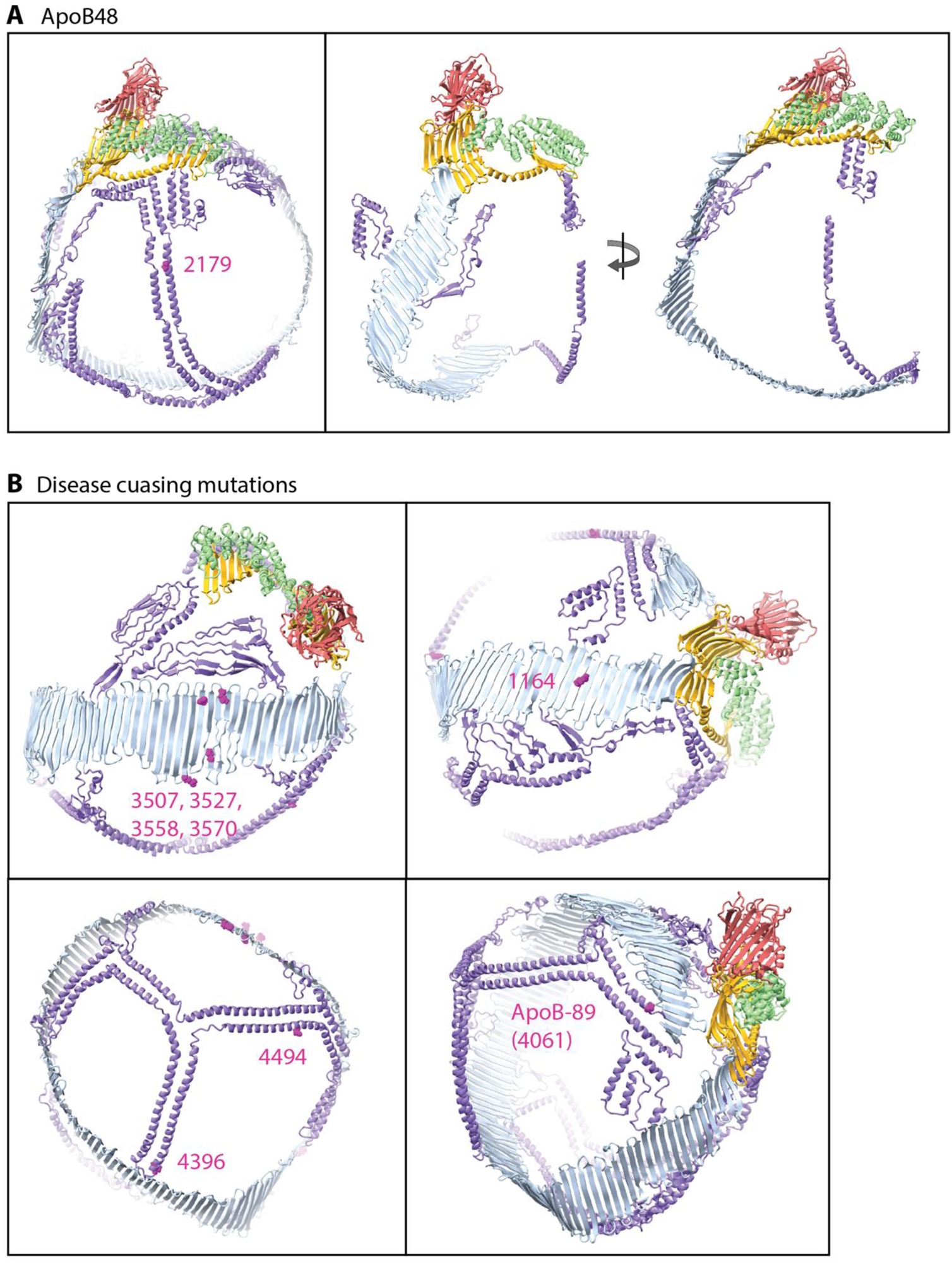
Modeling of apoB48 and mapping apoB100 disease-causing mutations. **A.** The structure of apoB100 with the C-terminal residue of apoB48 (2179) highlighted magenta (left) and with the C-terminus removed (right). **B.** Common and rare disease-causing mutations and truncations (apoB89) in apoB100.

## Notes

https://zenodo.org/doi/10.5281/zenodo.10723756

https://github.com/ZTBioPhysics/morphological-analysis-of-LDL-cryo-EM-2D-class-averages.git

### Competing Interest Statement

The authors have declared no competing interest.

