## Supplemental Information for "The Structure of ApoB100 from Human Low-density Lipoprotein"

#### Chemical crosslinking corroborates the proposed model of the C-terminal inserts.

A comprehensive chemical crosslinking study of apoB100 from human LDL was recently published that presented evidence of numerous long-range (in sequence) interactions [1]. The authors employed disuccinimidyl sulfoxide (DSSO), which is a mass spectrometry cleavable cross-linker that produces Cα–Cα distances between cross-linked lysine residues of ~26 Å. We sought to compare our structure with the reported long-range (>100 residues) cross-links, focusing on the C-terminal inserts where our cryo-EM map resolution is the lowest. Overall, we found that some of the reported distances match our structure exceptionally well (with a few angstroms), while some match reasonably well but still fall within the 26 Å limit, and others show poor agreement and exceed the 26 Å limit (Extended Data Figure 6A). However, most of the large discrepancies can be readily explained by considering the dynamic nature of the C-terminal inserts and the fact that our structure only captures one particle size (~20 nm diameter) compared to the full range of sizes captured in the cross-linking experiments. Looking at the R particle face, within arm 1 of insert 4, the authors found crosslinks between paired helices at the base (residue 2100 with residues 2387 and 2402) and tip (residue 2224 with 2270), which support its predicted topology (Extended Data Figure 6A). Most notably, a crosslink was observed between the tip of arm 1 (residue 2208) and insert 0 of the NTD (residue 766), which are within 10 Å in our model (Extended Data Figure 6A), supporting our proposed extended conformation stretching over 30 nm across the particle. An additional crosslink between the tip of arm1 (residue 2270) and insert 6 (residue 3210) was also reported, suggesting an alternative conformation. Although not readily supported by density in our cryo-EM map, such a conformation could conceivably be accessible if either insert 6 shifted to point away from the NTD, or if the particle was larger, and the β-belt was not as tightly wound. Finally, within arm 2 of insert 4, crosslinks were observed between the last helix of the descending edge (residue 2671) and the immediately adjacent β-belt segment (residues 936 and 942), supporting our overall placement of arm 2 (Extended Data Figure 6A).

Within the C-terminal inserts on the L-face of the particle, chemical crosslinks were detected between both sets of paired helices within arm 1 of insert 9 (residue 4076 with 4518 and residue 4132 with 4478), which support the overall trefoil topology (Extended Data Figure 6B). In addition, crosslinks were reported between the flexible linkers of arms 1 and 2 (residues 4103 and 4207) and inserts 7 (residue 3682) and 5 (residues 3148 and 3159), respectively, which support their localizations in our model (13Å and 25Å). An additional crosslink was detected between the flexible linker of arm 2 (residue 4207) and the edge of the β-belt just upstream of insert 3 (residue 1958) suggesting that arm2 may readily diffuse along the surface of the particle, a possibility supported by the relatively disordered density in this region of our cryo-EM map (Extended Data Figure 6B). Finally, one crosslink was observed between arm 3 (residue 4349) and the adjacent β-belt segment (residue 1696), although the residue in arm 3 was near the center of the trefoil rather than at the tip. Again, it’s conceivable that diffusion of arm 3 along the particle surface could enable such an interaction while not requiring an alteration of the overall topology of the trefoil, however, that possibility cannot be ruled out.

No long-range contacts were observed between the C-terminus of the β-belt and the NTD, between insert 8 and the NTD, or between insert 1 and either insert 9 or the C-terminus of the β-belt, despite these domains having exposed lysine residues within range of each other in our structure.

#### AlphaFold2 modeling of the LDL-r: insert 6 complex

As a preliminary experiment to test if insert 6 could be the LDL-r RBS, we used AF2 to model LDL-r: insert 6 complexes (Extended Data Figure 10A). The predicted interface spans nearly the entire fragment previously mentioned as the top candidate for the RBS, making contact with the positively charged lysine residues at the C-terminal end of the α-helix predicted to be important for LDL-r binding [2] (Extended Data Figure 10B-C). Alternative conformations of insert 6 were also predicted in which the α-helix formed two smaller helices that compacted together. It is difficult to see how these alternative conformations could form with insert 6 fully associated with the lipid membrane, however, given the poorly resolved cryo-EM density in this region, and previous results suggesting this domain is either not associated or only weakly associated with the particle surface, it is plausible that insert 6 could adopt alternative conformations not observed in our map.

#### ApoB100 N-linked glycosylation sites cluster near the putative RBS

ApoB100 contains 19 potential N-linked glycosylation sites (PNGS), 17 of which have been confirmed to harbor glycan modifications by mass-spectrometry and consist of a mixture of high-mannose and complex-type glycans [3, 4]. By mapping these PNGS to the apoB100 structure (Supplemental Figure 1) we find that 2 occur within the NTD (residues 185 and 983), 11 within the β-belt (1368, 1377, 1523, 2779, 2982, 3101, 3336, 3358, 3411, 3465, and 3895), 1 within insert 4, 1 within insert 6 (3224), and 2 within insert 9 (4237 and 4431). Of the PNGS that do not harbor glycan modifications, 1 falls within the unstructured N-terminal peptide (residue 34) and 1 within insert 4 (2560). Interestingly, 6 PNGS fall within residues 3000-3500 near the putative LDL-r binding region. This region has also been shown to be involved in proteoglycan binding as well, so these N-linked glycans may be important for cell attachment [5]. The PNGS at residue 3224 within insert 6 is located at the tip of the α-helix within the peptide fragment identified as the most probably LDL-r binding site [1]. The impact of this glycan modification on potential LDL-r binding is unknown.

### Supplemental Figures

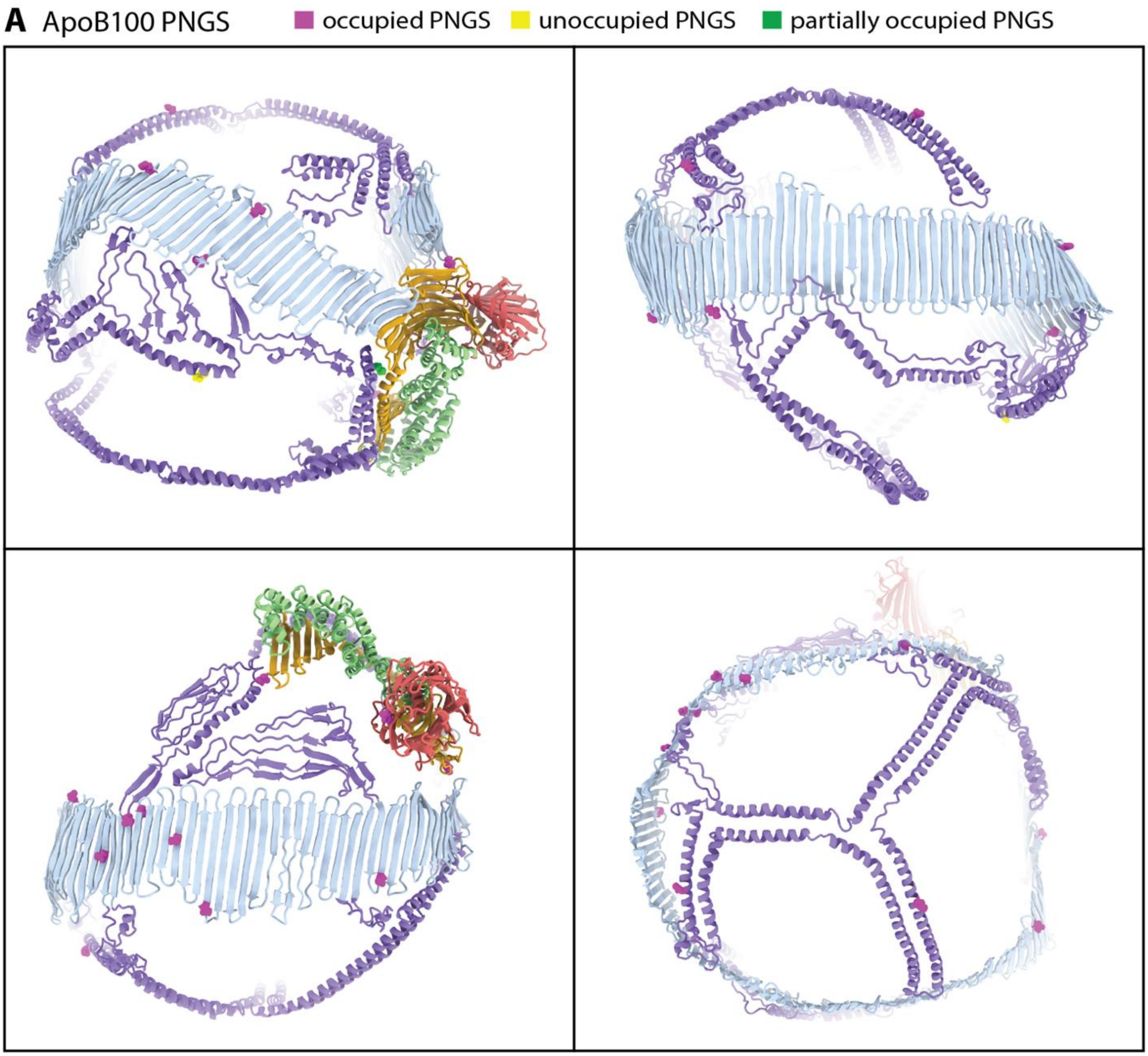


#### Supplemental Figure 1 | Mapping apoB100 PNGS.

**A.** Occupied, partially occupied, and unoccupied PNGS mapped onto the apoB100 structure and colored magenta.
